# CD8ɑ+ cells suppress SIV replication without evidence of viral immune escape during post treatment control

**DOI:** 10.1101/2025.11.18.689043

**Authors:** Ryan V. Moriarty, Olivia E. Harwood, Ethan P. Johnson, William Gardner, Corina C. Valencia, Andrew Conchas, Taina T. Immonen, Matthew R. Reynolds, Brandon F. Keele, Shelby L. O’Connor

**Author notes:** Address correspondence to Shelby L. O’Connor. Department of Cellular and Developmental Biology, Northwestern University, Chicago, IL.

## Abstract

Post-treatment control (PTC) is a rare phenomenon in which people living with HIV (PLWH) maintain viral control following ART interruption. Characterizing virus populations present in PTCs may help elucidate mechanisms of immunologic control, but this is challenging without detectable plasma viremia. To model PTC, eight Mauritian cynomolgus macaques (MCM) were infected with barcoded SIVmac239M and began an 8-month ART regimen two weeks post-infection (wpi). Six months following ART interruption, all MCM were rechallenged with non-barcoded SIVmac239 followed by CD8ɑ+ cell depletion two months later. Animals were grouped as viremic (n=5) or aviremic (n=3) based on the detection of plasma viremia between ART interruption and CD8ɑ+ cell depletion; all animals became viremic post-depletion. Barcode sequencing of plasma virus revealed that lineages with high pre-ART viral loads dominated the rebounding populations post-depletion and detectable rechallenge virus in two animals post-depletion. Additional sequencing of three CD8+ T cell epitopes within plasma viruses identified point mutations only in viruses isolated from the viremic group post-depletion. A second cohort of 5 MCM who initiated ART 8 wpi was examined to identify the impact of the timing of ART initiation on viral epitope diversity and showed increased diversity prior to ART initiation and following ART interruption. These results suggest that early ART initiation is associated with reduced diversity within cytotoxic T lymphocyte (CTL) epitopes and a longer time to rebound, as well as highlight an important role of restricting the emergence of CTL immune escape variants in increasing the likelihood of PTC.

**Importance:** While rare, a subset of PLWH, termed post-treatment controllers (PTCs), can maintain viral control following ART interruption. However, little is known about whether this control reflects a complete absence of viral replication or continual, subclinical replication. Here we address a key knowledge gap regarding how viral populations change during CD8ɑ+ cell-mediated PTC of SIV. We utilized our Mauritian cynomolgus macaque model of HIV infection, in combination with the barcoded SIVmac239M and deep sequencing, to characterize viral lineages and MHC-I-restricted CD8+ T cell epitopes throughout the study. Our findings demonstrate that early ART initiation limits viral diversity, that pre-ART replication predicts post-ART reactivation, and that CD8ɑ+ cells can suppress viral replication without evidence of immune escape. These insights establish MCMs as a valuable model for dissecting mechanisms of durable ART-free viral control and highlight the potential of CD8-mediated immune control as a therapeutic target for HIV cure strategies.

## Introduction

While ART is required for most people living with HIV (PLWH) to maintain viral control, rare individuals, termed post-treatment controllers (PTCs), can maintain undetectable viremia for months to years following ART interruption (1–4). Unfortunately, the exceptionally low frequency (approximately 4-16%) of these individuals in the population (3,5), combined with their low or undetectable viremia, has hindered comprehensive analyses of viral population diversity, replication dynamics, and reservoir stability between PTCs and ART-treated individuals. Studies of durably suppressed, ART-treated individuals report virtually no evidence of viral evolution in proviral DNA sequences sampled during treatment or in plasma viruses isolated at the time of ART interruption (6–9), suggesting efficient inhibition of viral replication. When examining the sequence of replication-competent virus during transient viremia or following ART interruption, the most commonly identified viruses in the rebounding population were typically those present at the highest frequency at the time of therapy initiation (10–13), suggesting little change in the composition of the reservoir during ART. Due to the limited data available for PTCs, it is not known whether the composition of the replication-competent viral reservoir is stagnant during the time when HIV is controlled after stopping ART, or if HIV is replicating at levels below the limit of detection, leading to increased within-host viral diversity and the potential emergence of immune escape mutations.

Cytotoxic CD8+ T lymphocytes (CTLs) play a critical role in the control of HIV/Simian immunodeficiency virus (SIV) infection and are a major driver of viral evolution (14,15). High frequencies of potent, virus-specific CTL responses have been associated with post-peak viral load decline and viral suppression during acute infection (16,17). In response to this CTL-mediated selective pressure, immune escape variants within MHC-I restricted CD8+ T cell epitopes can emerge (14,17–19). Although the emergence of CTL escape variants has been associated with loss of viral control (20,21), CTL escape variants have also been identified in individuals that naturally maintain control of viremia or subsequently re-control viremia following transient loss of control (18,22,23). However, it is not understood how PTCs may differ from spontaneous control of untreated viremia and if mutations accumulate within CTL epitopes of viruses found in PTCs during these periods of viral control in the absence of continual ART.

SIVmac239M, a molecularly barcoded, clonal SIV strain, has emerged as a vital tool to examine distinct SIV lineages experimentally. SIVmac239M differs from the clonal SIVmac239 by a 34bp insertion between *vpx* and *vpr* that allows for the distinction of approximately 10,000 unique viral lineages using next-generation sequencing (24). This unique viral strain has been used to examine the number and identity of rebounding lineages following ART interruption (24,25), viral reactivation rates (26–29), and the impact of route of infection on the number of viral lineages present during acute infection (30). Using this molecularly barcoded viral strain, we can examine the persistence of distinct SIVmac239M lineages within an animal over time and how the composition of replication-competent viral lineages within the viral reservoir may change during PTC.

We recently reported that Mauritian cynomolgus macaques (MCMs) that began ART at two weeks post-SIV infection could maintain control of viremia for at least six months after ART interruption (31,32). Here, we sought to understand the viral populations and the number and composition of unique lineages persisting over time and how this may influence the likelihood of establishing PTC in these animals. We utilized deep sequencing to characterize the SIV lineages present and MHC-restricted CD8+ T cell epitopes in viruses isolated from MCM that were infected with SIVmac239M, underwent an 8-month ART regimen beginning 2 weeks post-infection (wpi), were rechallenged with the isogenic, non-barcoded SIVmac239 six months after ART interruption, and given a CD8ɑ+ cell depleting antibody 8 weeks after rechallenge. By sequencing the molecular barcode throughout the study, we quantified the number of unique lineages that established persistent infection and determined whether our experimental interventions impacted the composition of viral lineages that could be detected in the plasma as well as determine susceptibility to isogenic rechallenge. To further evaluate the effects of early versus late ART initiation on the composition of the viral population, we included a second cohort of MCM infected with SIVmac239 who initiated ART at 8 wpi. We compared the sequence diversity of 3 MHC class I-restricted viral epitopes in viruses isolated from the two cohorts just prior to ART initiation and when SIV became detectable after stopping ART. The results of our study highlight the potential benefits of earlier ART initiation and antiviral CD8+ cells in restricting viral diversity and enhancing the likelihood of PTC, as well as support MCMs as a potential model of PTC.

## Results

### Description of Early ART study from which samples were derived

The study timeline and plasma viral loads for each animal are depicted in Figure 1A and B, respectively. Briefly, 8 MCM lacking the M1 MHC haplotype associated with spontaneous SIV control (33) were infected intravenously with 10,000 infectious units (IU) of SIVmac239M and began receiving daily antiretroviral treatment (ART) consisting of dolutegravir, tenofovir disoproxil fumarate, and emtricitabine at 14 days post-infection. Plasma viral loads show high pre-ART viremia, peaking between 11- and 14-days post-infection, followed by rapid viral suppression once ART was initiated, with no detectable viremia during the 8-month ART regimen. Animals were monitored for viral rebound for seven months, during which time 4 of 8 animals had detectable, often transient, viremia. At this point, all animals were rechallenged intravenously with 100 TCID_50_ non-barcoded SIVmac239, after which one additional animal became viremic. Two months following isogenic rechallenge, all animals were depleted of CD8ɑ+ cells, which led to high levels of viremia in all 8 MCMs. All animals were necropsied approximately two months after depletion. The animals with no detectable viremia between ART interruption and CD8ɑ+ depletion are hereafter referred to as “aviremic” (n=3; Figure 1B, blue). The remaining animals with detectable viremia above 10^3^ copies/mL of plasma for at least two consecutive time points between ART interruption and CD8ɑ+ depletion are referred to as the “viremic” cohort (n=5; Figure 1B, black).

**Figure 1.**
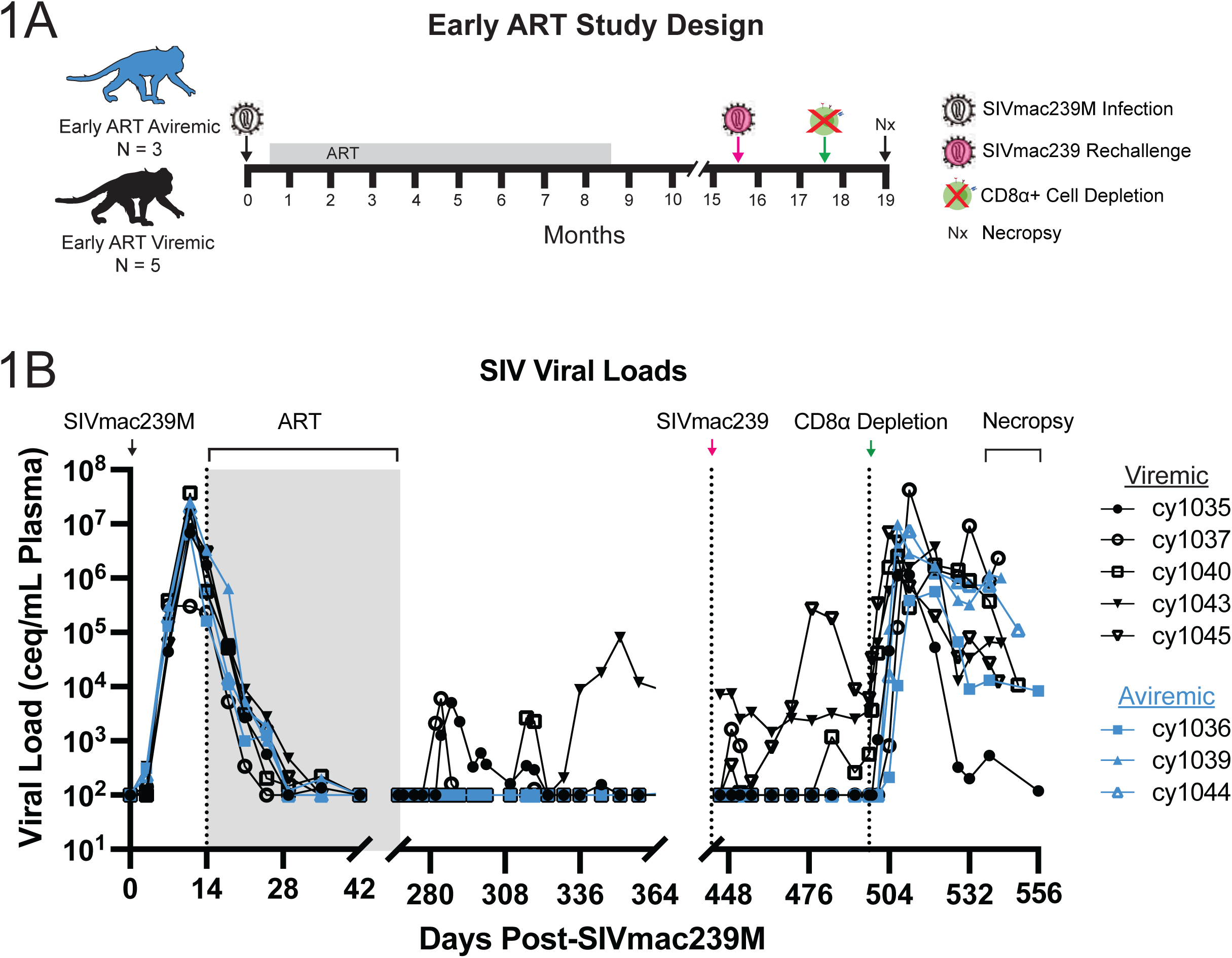
(A) Study design and timeline. Intravenous SIVmac239M infection is indicated by the white virion, ART administration is indicated by the grey bar, intravenous SIVmac239 rechallenge is indicated by the pink virion, CD8ɑ+ cell depletion is indicated by the green x-ed out cell, and necropsy is indicated by “Nx.” Animals with no detectable viremia between ART interruption and CD8ɑ+ cell depletion are referred to as “Early ART aviremic” (n=3, blue). The remaining animals with detectable viremia above 10^3^ copies/mL of plasma for at least two consecutive time points between ART interruption and CD8ɑ+ cell depletion are referred to as the “Early ART viremic” (n=5, black). (B) SIV viral loads over the course of the animal study, with each animal indicated by a unique color and symbol combination, with blue representing aviremic animals and black representing viremic animals. ART was administered between days 14 and 268 post-SIVmac239M infection and indicated by the grey shaded area. Intravenous SIVmac239 rechallenge occurred on day 442 post-SIVmac239M infection. Administration of a CD8ɑ+ cell-depleting antibody occurred on day 497 post-SIVma239M infection. Necropsies were conducted between days 539 and 556 post-SIVmac239M infection.

### The number of viral lineages during acute infection was not associated with the detection of transient plasma viremia following ART interruption

We first evaluated if the number of SIVmac239M lineages detectable in the plasma prior to ART initiation was associated with whether an animal rebounded after ART interruption. We sequenced the molecularly barcoded SIVmac239M isolated from plasma and evaluated the total number of unique lineages present in each animal prior to ART initiation and following CD8ɑ+ cell depletion (Figure 2). We found no difference in the number of detectable lineages at peak, pre-ART plasma between MCMs that rebounded off-ART and those that did not. However, the aviremic animals had a significantly greater number of distinct lineages when compared to the viremic animals following CD8ɑ+ cell depletion (p = 0.0357, Mann-Whitney test). These results suggest that while a similar number of lineages may be present during the first two weeks following intravenous challenge, there may be an association with viral control following ART interruption and the number of viral lineages that are able to rebound following CD8ɑ+ cell depletion.

**Figure 2.**
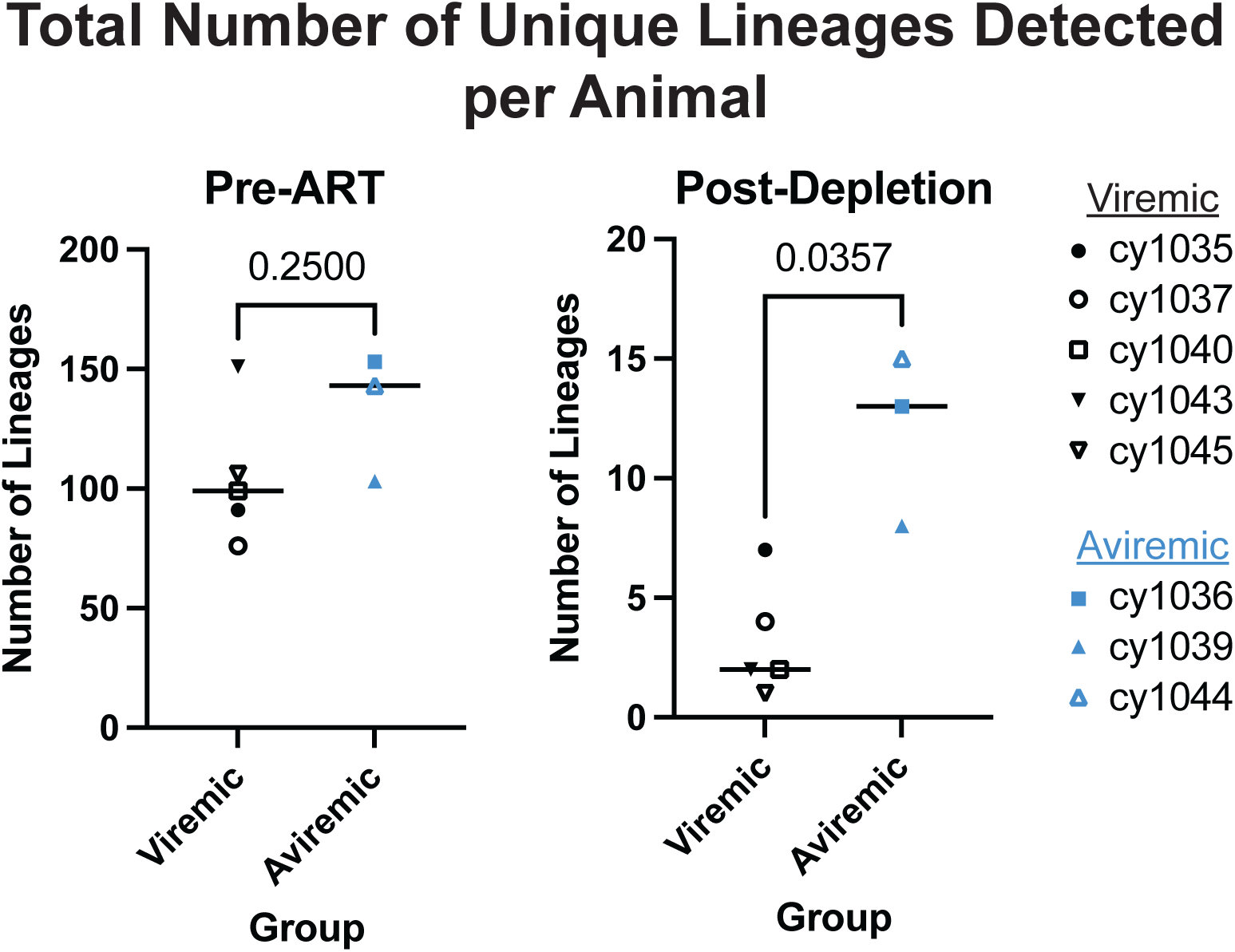
Number of unique SIVmac239M lineages detected in the plasma during the first two weeks of SIVmac239M infection, prior to ART initiation (A) and following CD8ɑ+ cell depletion (B). Animals are coded by unique color and symbol combinations, with blue representing aviremic animals and black representing viremic animals. Only barcodes present at a frequency of 0.002 or greater, which corresponds to 1/minimum input templates of time points included in this analysis, were included. Significance is determined by Mann-Whitney test.

### Few viral lineages reactivate following ART interruption

After approximately eight months of daily ART administration, ART was interrupted and plasma was monitored for seven months to detect viral rebound. Three of 8 animals (cy1035, cy1037, and cy1040) had transiently detectable viremia and one additional animal (cy1043) had a delayed rebound that was sustained for the rest of the study (Figure 1B). To evaluate the composition of viral lineages present in the rebounding viral populations, we sequenced the rebounding virus with viral loads greater than 10^3^ copies/mL of plasma in these four animals. We identified between 2 and 4 unique lineages from each time point in each animal (Supplemental Figure 1A-D), highlighting the strong genetic bottleneck between primary infection (pre-ART) and rebound following ART interruption.

### Most animals were not susceptible to rechallenge with SIVmac239

We next determined if these animals were susceptible to infection with a homologous challenge virus. Approximately six months following ART interruption, all animals were intravenously rechallenged with 100 TCID_50_ of the non-barcoded SIVmac239. Rechallenge virus, as identified by the detection of SIV lacking a barcode between *vpx* and *vpr*, was detectable in the plasma immediately following challenge in all animals at a frequency between 94 and 100% (Supplemental Figure 2), confirming that the rechallenge virus was successfully administered to each animal. To determine if the rechallenge virus successfully established infection, we sequenced the molecular barcode region of plasma viruses from time points with a viral load of 10^3^ copies/mL of plasma or greater from the day after rechallenge until necropsy (Supplemental Figure 1A-H). Interestingly, the rechallenge virus was only detected in one animal (cy1037) immediately following challenge and one additional animal (cy1044) following CD8ɑ+ cell depletion (Figure 3 and Supplemental Figure 1B, H). In the case of cy1037, we found that the majority of the detectable virus population post-rechallenge and post-depletion consisted of the rechallenge virus, with only a small proportion (up to 12%) consisting of one other lineage (Figure 3A and Supplemental Figure 1B). In the case of the aviremic animal, cy1044, the rechallenge virus was found only at one time point post-rechallenge, approximately seven days post-CD8ɑ+ cell depletion at a frequency of approximately 1% (Figure 3B and Supplemental Figure 1H), suggesting that while the rechallenge virus was able to establish infection and contribute to the post-depletion viral population in these two animals, this was limited, replication was transient, and the lineage could be spontaneously controlled.

**Figure 3.**
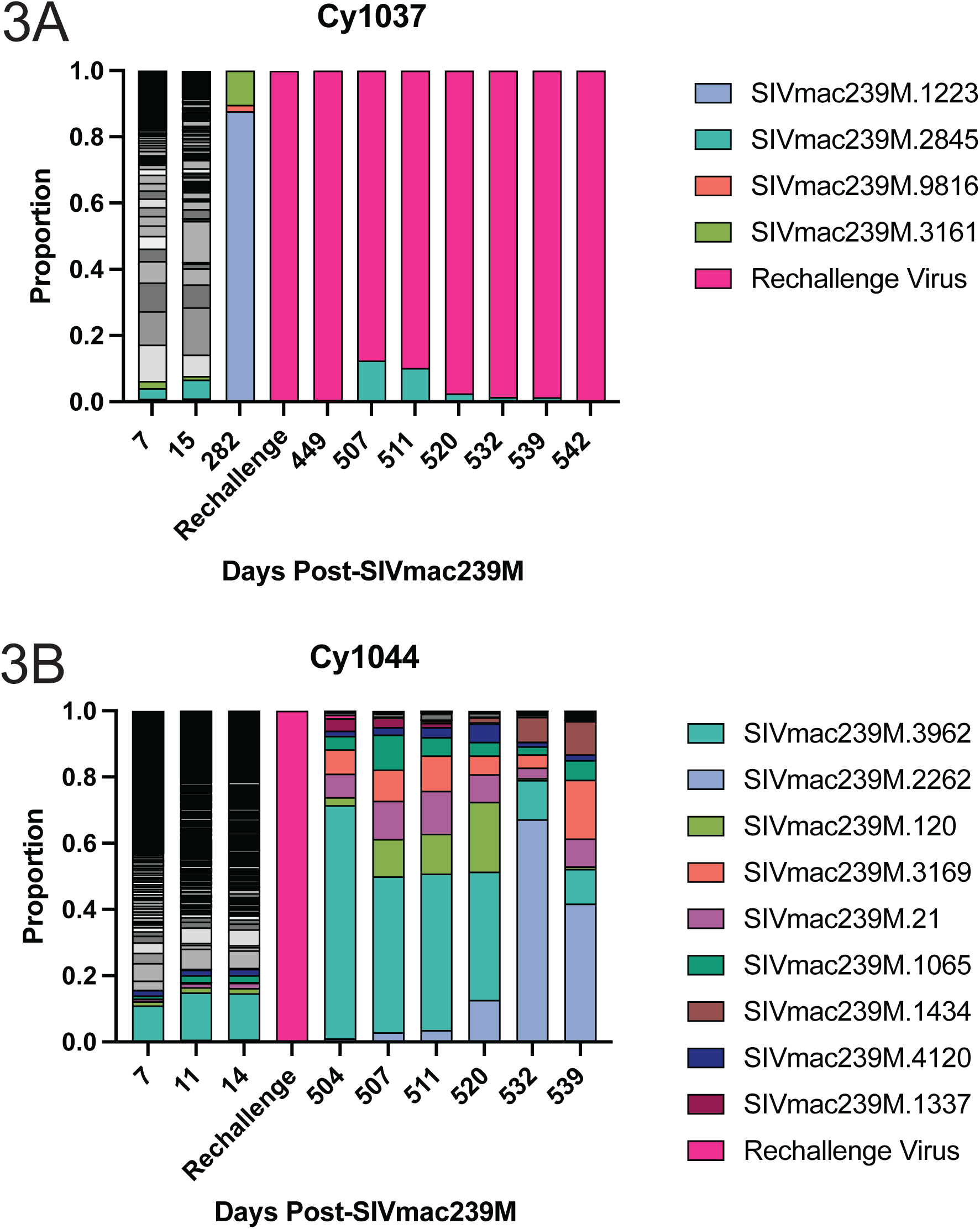
Distribution of SIVmac239M lineages throughout time for the viremic (A, cy1037) and aviremic (B, cy1044) animals which had detectable rechallenge virus following intravenous rechallenge. Each unique lineage detected post-ART or post-depletion is indicated by color, with the rechallenge virus indicated in bright pink. In the case of cy1044 (B), only the top ten lineages detected post-depletion are shown due to color and space constraints; the remaining lineages are show in shades of grey.

### Viral lineages detected off-ART typically originate from the highest-replicating lineages pre-ART

We next compared the viral lineages detected pre-ART to all the lineages detected after ART interruption (including after CD8ɑ+ cell depletion). We hypothesized that lineages that reactivated following ART interruption would have had the highest extent of viral replication pre-ART, similar to what was seen previously (25). Each lineage was therefore ranked by viral load at peak viremia (between days 11 and 14 post-infection), with the rechallenge virus last when present. Viral load for each lineage during peak pre-ART viremia is shown in grey and represents the baseline for each barcode lineage, whereas cumulative viral loads for each lineage during each study phase is shown in different colors stemming from the initial peak pre-ART symbol (Figure 4).

**Figure 4.**
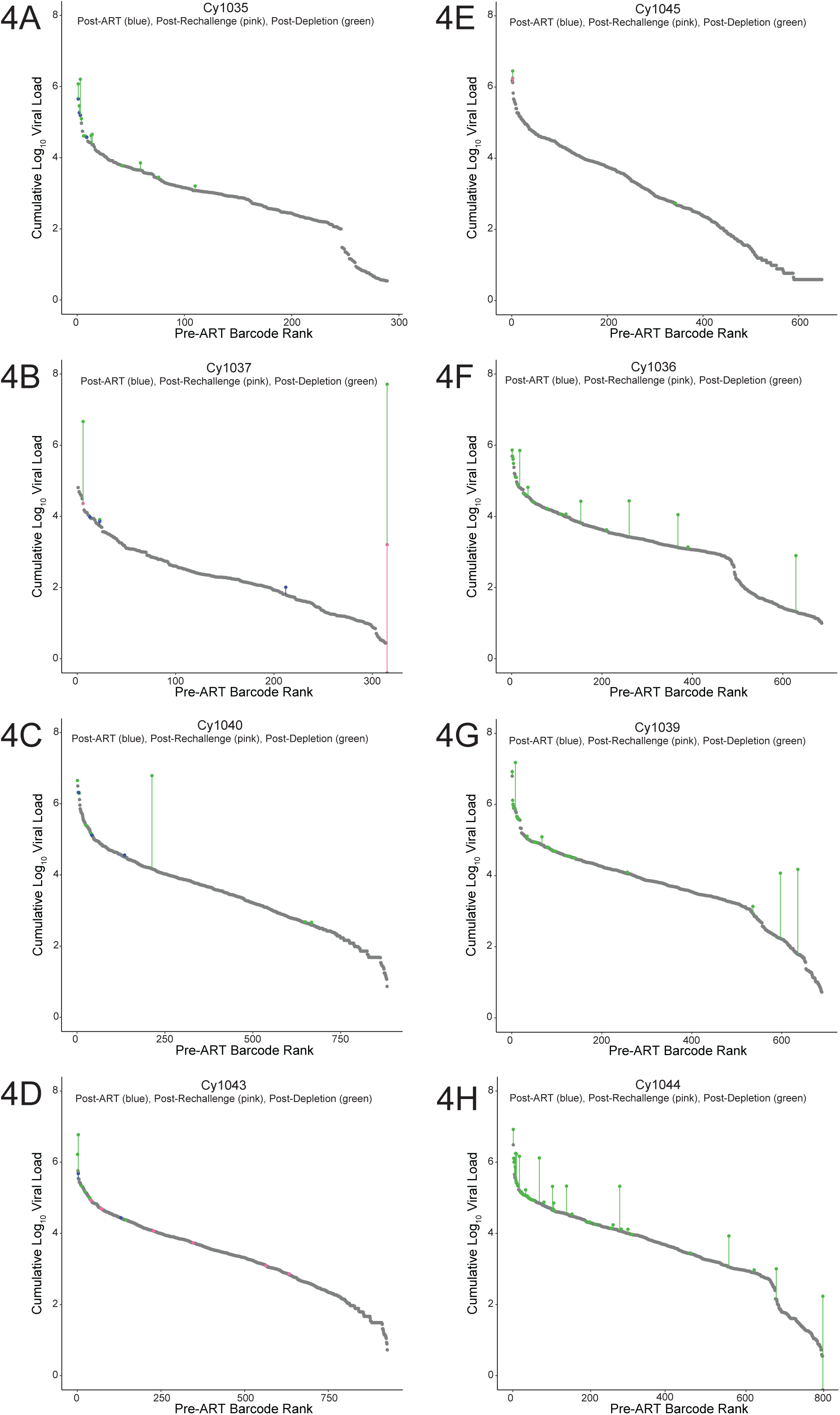
Detection of SIVmac239M lineages following ART interruption (blue), SIVmac239 rechallenge (pink) and after CD8a+ cell depletion (green). Grey dots represent the peak pre-ART viral load of each lineage, with lineages ranked from highest pre-ART replication to lowest pre-ART replication. The rechallenge virus is present on the far right of graphs B (cy1037) and H (cy1044), as this lineage was absent prior to ART initiation but was detected post-rechallenge and depletion.

When examining the barcode sequences of the viremic animals, we found that viral lineages detected in rebounding viral populations typically originated from the top of the pre-ART distribution, with lineages replicating to a high extent in pre-ART (grey), post-ART (blue), and post-rechallenge (pink) also replicating post-depletion (green) population. However, we found that not all lineages present in the post-depletion population also reactivated post-ART or post-rechallenge, such as in animals cy1037 (Figure 4B) and cy1040 (Figure 4C). These results suggest that rebound virus can originate from multiple, distinct reservoirs, and that prior replication during a treatment interruption is not required, or necessarily predictive, of viral replication post-depletion.

Following CD8ɑ+ cell depletion, all animals became viremic. We found that the rebounding lineages post-depletion in previously aviremic animals were distributed widely throughout the peak pre-ART barcode distribution, whereas post-depletion rebounding lineages in viremic animals were predominantly present in the top of the barcode distribution (green symbols, Figure 4F-H). This result is consistent with previous studies conducted in rhesus macaques who have undergone analytical treatment interruptions both with and without CD8ɑ+ cell depletion (25), supporting the hypothesis that while lineages with high pre-ART viral replication are able to seed a large portion of the viral reservoir and rebound following treatment interruption, even lineages with low pre-ART viral replication contribute to the replication-competent viral reservoir, particularly when CD8-mediated control of viral replication is removed.

### Pre-ART viral replication predicts likelihood of post-ART and post-depletion lineage reactivation

Following HIV/SIV infection, only a subset of viral lineages establish productive infection, comprise the replication-competent viral reservoir, and rebound following treatment interruption (24,29,34,35). However, it is unclear whether the composition of the replication-competent viral reservoir changes during periods of off-ART replication. We hypothesized that the pre-ART viral load area under the curve (AUC), representing its cumulative contribution to the viral reservoir prior to ART initiation, would be a significant predictor of whether that lineage was detected following ART interruption, but prior to isogenic viral rechallenge, suggesting a direct relationship between the size of the lineage-specific viral reservoir and the likelihood of that lineage rebounding post-ART. We further hypothesized that viral replication of a given lineage during periods of transient viremia, either post-ART or post-depletion, would contribute to viral reservoir reseeding and increase the likelihood that the replicating lineage would also be detected following CD8ɑ+ cell depletion.

Consistent with our hypothesis, we found that pre-ART viral load AUC was a significant predictor of reactivation in 3 of 4 viremic animals, with animal cy1037 being the exception (Table 1, Model 1, p <0.05). When we extended our analysis to evaluate whether the initial reservoir size was a significant predictor of reactivation following ART until necropsy, we found this was the case in both viremic and aviremic animals, except for cy1037 and cy1045 (Table 1, Model 2, p < 0.05).

**Table 1.** Summary of logistic regression model parameters. Model 1 examines the virus population in the plasma following ART interruption but prior to isogenic viral rechallenge (between days 268 and 449 post-SIVmac239M infection). Model 2 examines the plasma virus population at all time points following ART interruption. The sole explanatory variable for Model 1 and Model 2 is the area under the curve (AUC) of a given lineage prior to ART initiation. Models 3 and 4 incorporate both the AUC of a given lineage prior to ART initiation and a binary indicator variable for whether a lineage was detected between ART interruption and isogenic rechallenge (Model 3) or between isogenic rechallenge and CD8ɑ+cell depletion (Model 4).

|  |  |  | cy1035 | cy1037 | cy1040 | cy1043 | cy1045 |  | cy1036 | cy1039 | cy1044 |
| --- | --- | --- | --- | --- | --- | --- | --- | --- | --- | --- | --- |
| Post-ART | Model 1:<br>Does pre-ART VL AUC predict post-ART rebound? | AIC | 13 | 34 | 34 | 26 | - |  | - | - | - |
|  |  | Intercept (p-value) | -36.9 (0.0125) | -7.05 (5.58e-04) | -14.4 (2.33e-04) | -17.4 (1.18e-04) | - |  | - | - | - |
|  |  | Pre-ART VL AUC (p-value) | 6.71 (0.0155) | 0.831 (0.111) | 1.8 (7.16e-03) | 2.36 (0.0298) | - |  | - | - | - |
| Post-Depletion | Model 2:<br>Does pre-ART VL AUC predict if a lineage will be detected at any point post-ART interruption? | AIC | 61 | 52 | 103 | 117 | 28 |  | 161 | 200 | 300 |
|  |  | Intercept (p-value) | -15.8 (4.05e-07) | -5.45 (4.88e-07) | -9.46 (4.52e-08) | -11 (2.64e-08) | -9.72 (8.05e-03) |  | -7 (2.4e-09) | -9.43 (9.79e-12) | -9.84 (2.45e-15) |
|  |  | Pre-ART VL AUC (p-value) | 3.03 (5.35e-06) | 0.525 (0.0843) | 1.14 (4.66e-04) | 1.52 (9.61e-05) | 0.977 (0.187) |  | 0.922 (3.88e-04) | 1.35 (4.98e-07) | 1.55 (2.86e-10) |
|  | Model 3:<br>Does detecting a lineage between ART interruption and isogenic rechallenge improve the likelihood of detecting that lineage post-depletion? | AIC | 60 | 33 | 86 | 44 | - |  | - | - | - |
|  |  | Intercept (p-value) | -14 (1.47e-05) | -5.71 (5.66e-06) | -8.63 (2.33e-06) | -22.4 (3e-05) | - |  | - | - | - |
|  |  | Pre-depletion VL AUC (p-value) | 2.62 (2.21e-04) | 0.304 (0.434) | 0.917 (0.01) | 3.45 (2.78e-04) | - |  | - | - | - |
|  |  | Post-ART Detection (p-value) | 17.1 (0.993) | 3.85 (0.0103) | -13.5 (0.992) | 3.44 (0.109) | - |  | - | - | - |
| | Model 4:<br>Does detecting a lineage between isogenic rechallenge and CD8α+ cell depletion improve the likelihood of detecting that lineage post-depletion? | AIC | - | 17 | - | 47 | 21 | | - | - | - |
|  |  | Intercept (p-value) | - | -11.1 (0.058) | - | -23.2 (2.34e-05) | -6.84 (2.64e-04) |  | - | - | - |
|  |  | Pre-depletion VL AUC (p-value) | - | 1.56 (0.244) | - | 3.61 (1.82e-04) | 0.128 (0.801) |  | - | - | - |
|  |  | Post-Rechallenge Detection (p-value) | - | 28.7 (0.994) | - | -13.3 (0.995) | 23.6 (0.995) |  | - | - | - |

When evaluating if replication of a given lineage post-ART (Model 3) or post-rechallenge (Model 4) improved the model, we found that although pre-ART viral load AUC remained a significant predictor for most viremic animals, the inclusion of these variables did not typically improve the model fit, with most comparisons showing similar AIC values (Models 3 and 4 compared to Model 2). Additionally, post-ART or post-rechallenge detection was not a significant predictor of rebound post-depletion in any animal (Models 3 and 4, p > 0.1), except for cy1037 (Model 3, p = 0.01), whose viral population consisted of primarily the rechallenge virus following isogenic rechallenge (Figure 3 and Supplemental Figure 1B). Together, these results suggest minimal influence of post-ART and post-rechallenge detection on the post-depletion population, but do not definitively support or refute the continual reseeding hypothesis (25), likely due to the low level of viral replication that occurred between ART interruption and CD8ɑ+ cell depletion.

### CTL epitope sequences present in viruses from the aviremic group matched the inoculum at the time of CD8ɑ+ cell depletion, but accumulated variants as the CD8ɑ+ cells returned

The observed viral rebound associated with depletion of CD8ɑ+ cells indicated that CD8ɑ+ cells were responsible for PTC (32). To determine whether the existing CD8ɑ+ cells had selected for viral variants during PTC, we characterized the sequences of three MHC class I-restricted CD8+ T cell epitopes from viruses isolated from seven animals at the time of CD8ɑ+ cell depletion to determine if they had accumulated mutations. Animal cy1037 was excluded from epitope analysis because nearly 100% of the viral population could be attributed to the rechallenge, which prohibited the study of viral epitopes in this animal during periods of PTC.

All animals expressed at least one copy of the M3 MHC haplotype (Supplemental Table 1, (31,32)), which contains the *Mafa-A1*063* and *Mafa-B*075* MHC class I alleles. We focused on two epitopes restricted by Mafa-A1*063 (Gag_386-394_GW9 and Nef_103-111_RM9) and one epitope restricted by Mafa-B*075 (Rev_59-68_SP10) that commonly develop immune escape mutations during acute SIV infection (36). Prior to ART initiation, the CTL epitope sequences were nearly 100% identical to the inoculum, as expected (Figure 5A). Approximately one week after administering the CD8ɑ-depleting antibody, the three CTL epitope sequences present in viruses from animals in the Early ART aviremic group were predominantly wild-type, whereas those present in viruses from the Early ART viremic animals were almost entirely variant (Figure 5B). Approximately 3-4 weeks following CD8ɑ+ cell depletion, bulk, but not peptide-specific, CD8ɑ+ cells returned in the peripheral blood of most animals (Supplementary Figure 3). At this point, viral variants were detectable in the sequences of the targeted epitopes from all animals in both groups (Figure 5C). While we wanted to determine if unique barcoded SIV lineages emerged coincident with the detection of variant epitopes, the three epitopes we examined were too far away from the barcode (all greater than 2kb away from the barcode region) to perform a linkage analysis given our sequencing methodology.

**Figure 5.**
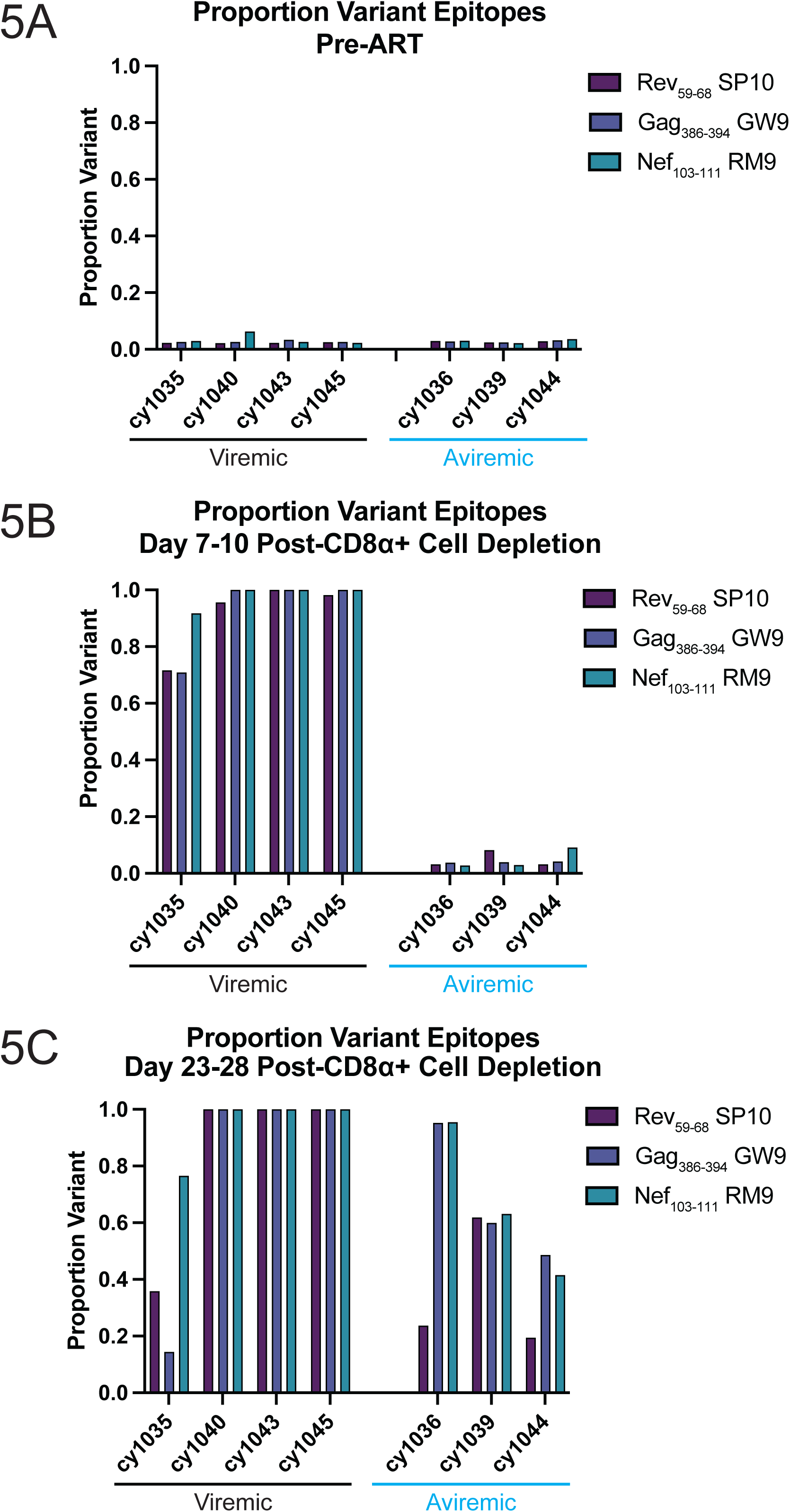
Proportion of variant Gag_386-398_GW9 (blue), Nef_103-111_RM9 (teal), and Rev_59-68_SP10 (purple) epitope sequences in viremic (left, black) and aviremic (right, blue) animals prior to ART initiation (A), 7-10 days following CD8ɑ+ cell depletion (B), and 21-28 days post-CD8ɑ+ cell depletion (C).

### Viral rebound in MCMs is associated with the presence of CTL variants present prior to ART initiation

While the relationship between CTL escape and PTC is unknown, the presence of CTL escape mutations has been associated with poor viral control in the absence of any ART (20,37,38). Therefore, we also examined five MCMs from the Harwood et. al. study that started a 22-month course of ART 8 weeks post-infection, four of which had at least one copy of the M3 MHC haplotype (Supplemental Table 1, (32)). These animals had detectable, sustained viremia (range 1.25 × 10^2^ to 4.85 × 10^5^ copies/mL) between two and six weeks after ART interruption (Late ART cohort, Figure 6, (32)), in stark contrast to the Early ART cohort (Figure 1B). We sequenced the plasma virus from the Late ART cohort and the Early ART viremic cohort immediately prior to ART initiation and at the first detection of virus greater than 10^3^ copies/mL of plasma after ART interruption. We calculated the composition of variant epitope sequences for the three MHC-restricted epitopes Gag_386-394_GW9, Rev_59-68_SP10, and Nef_103-111_RM9. These epitopes were primarily wild type at both time points for the Early ART viremic cohort (Figures 5A and 7). In contrast, all epitopes were largely variant at both time points in the Late ART cohort, even if the specific variants differed between time points (Figure 7).

**Figure 6.**
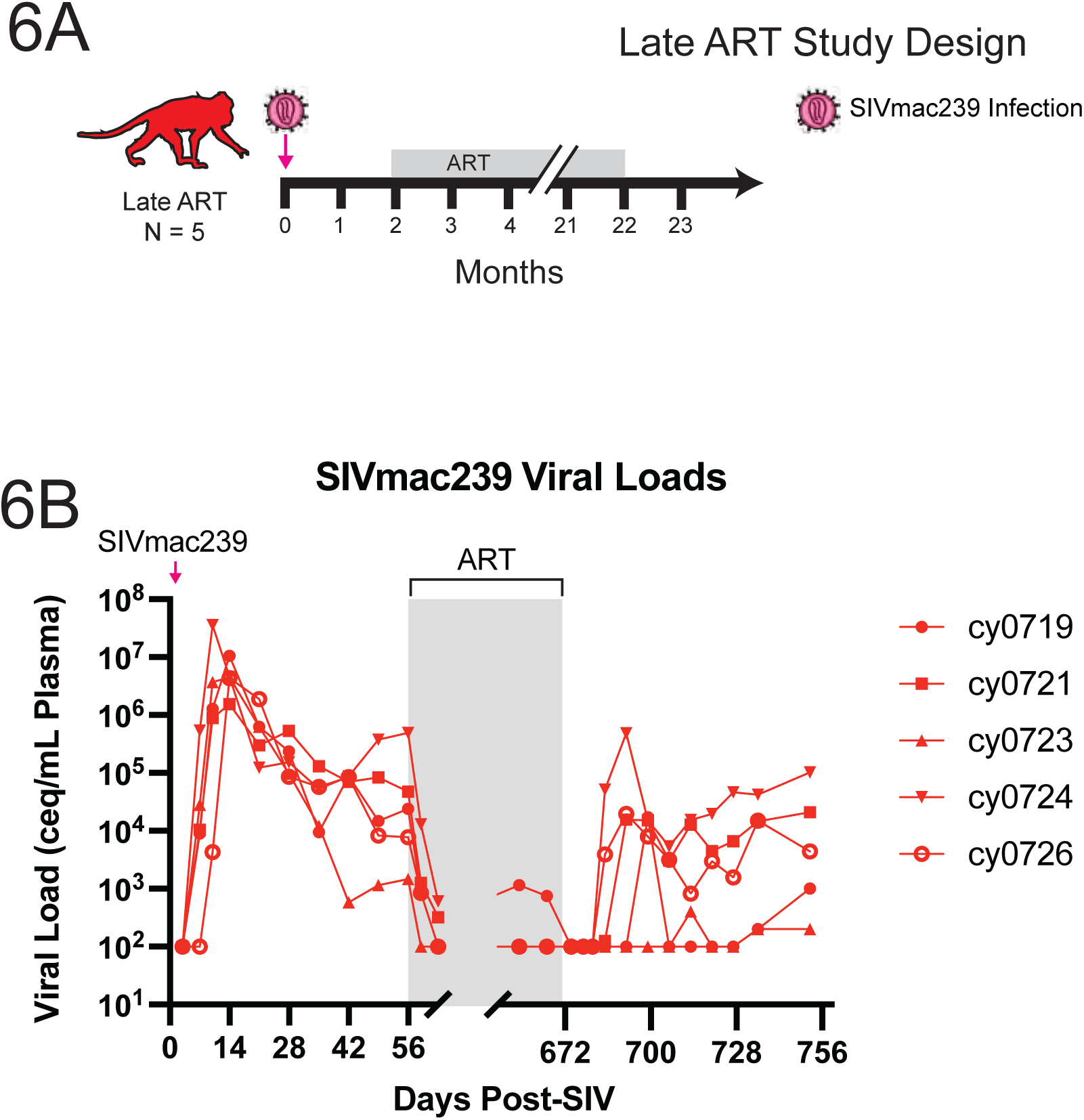
Study design and timeline for the Late ART animals (A). All 5 animals were infected intrarectally with SIVmac239 and began ART 8 weeks post-infection. ART was withdrawn after approximately 22 months. SIV plasma viral loads for the Late ART animals (B). ART administration is shown by the grey shaded bar. Animals are indicated by unique symbols.

**Figure 7.**
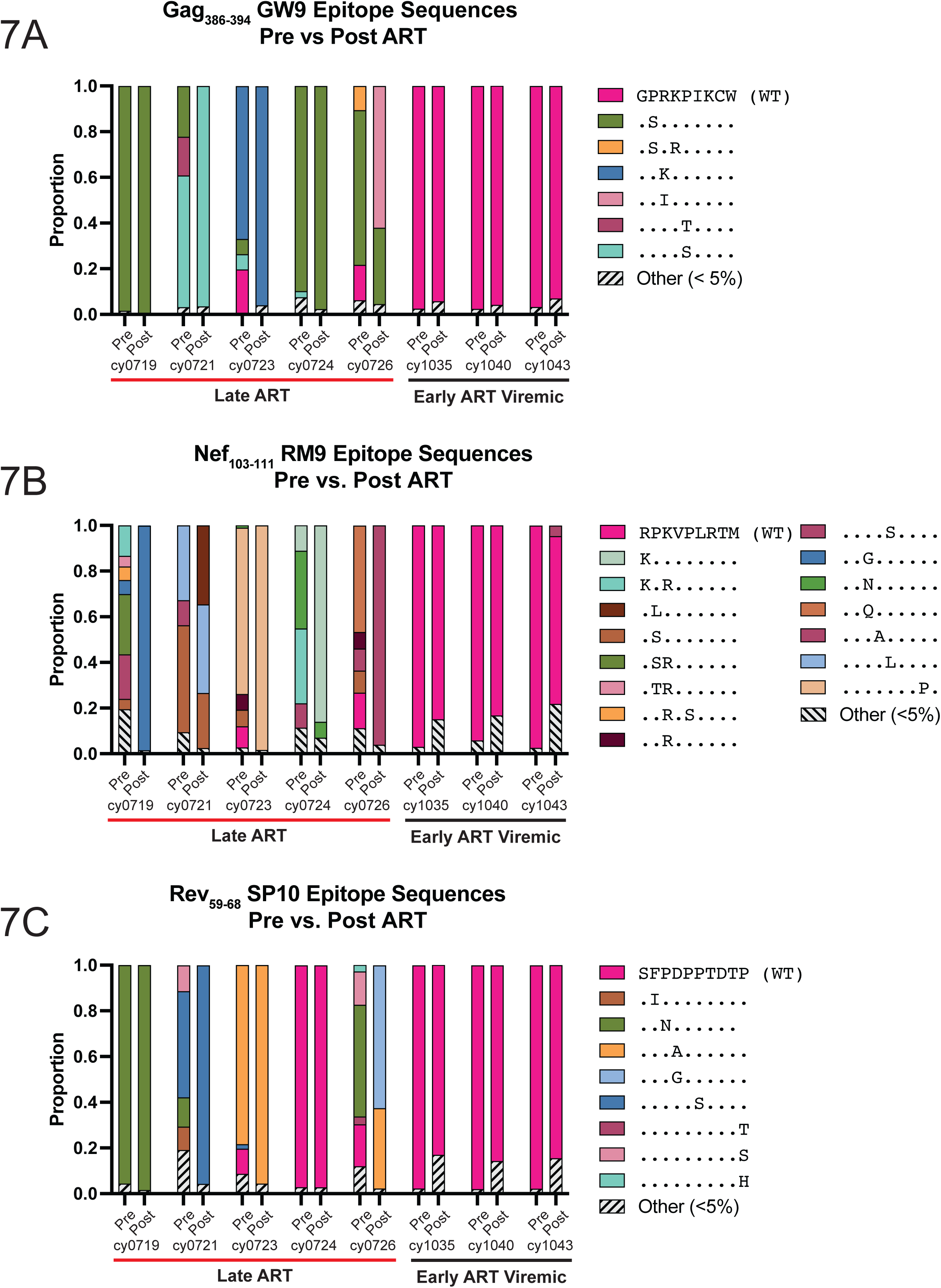
Gag_386-398_GW9 (A), Nef_103-111_RM9 (B), and Rev_59-68_SP10 (C) epitope sequences pre-and post-ART for Late ART (left, red) and Early ART viremic (right, black) animals. The “pre-ART” time point was between 11 and 14 days post-SIVmac239M infection for the Early ART cohort and 49 days post-SIVmac239 for the Late ART cohort. The “post-ART” time point was the first time point following ART interruption with a viral load greater than 10^3^ copies/mL. Wild type epitope sequences are in pink bars. Epitope sequences that comprised less than 5% of the total population are in the striped grey and black bars. Variant epitope sequences are shown in different colors. Residues matching the wild-type epitope sequences are indicated by dots and nonsynonymous residues are indicated by letters.

We calculated the Bray-Curtis dissimilarity index, which compares the compositional dissimilarity between two populations, for the three MHC-I-restricted T cell epitopes examined in each animal to determine the difference between pre-ART and post-ART populations. Across all three epitopes, we found that the Late ART cohort had a slightly higher Bray-Curtis value (median > 0.37) when compared to the Early ART viremic cohort (median < 0.16) (Supplemental Figure 4). While statistical significance was not reached, the greater dissimilarity between pre-ART and post-ART time points in the Late ART animals suggests that the presence of escape mutations prior to ART initiation may contribute to continual epitope evolution despite persistent ART treatment and the absence of detectable viremia.

## Discussion

In this study, we took advantage of a unique opportunity to examine viral populations in eight SIVmac239M-infected MCM who initiated ART at 2 weeks post-infection and then exhibited variable degrees of PTC, as described in Harwood et. al. (32). This allowed us to conclude that while most animals were not susceptible to infection by the rechallenge virus, the two animals that were susceptible differed greatly with respect to the extent of rechallenge virus detectable in the plasma. We also found that lineages rebounding at any point following ART interruption typically had high levels of pre-ART viral replication, and that pre-ART replication, but not post-ART or post-rechallenge replication, was predictive of viral reactivation. Finally, we discovered that the aviremic animals did not develop immune escape mutations in CD8+ T cell epitopes during the period of PTC. Cumulatively, this study highlights the importance of evaluating viral populations in PTCs, specifically with respect to how the composition of these viral populations may be influenced by transient viral replication and selective pressure exerted by host immune responses.

Due to the exceptional rarity of PTCs, examination of viral populations within these individuals has been limited and has primarily focused on the size and composition of the proviral reservoir (39,40). However, understanding if, and to what extent, viral replication and evolution is ongoing in these individuals has major implications for sustained viral control, as the development of immune escape mutations has been found to precede loss of viral control in elite controllers (21,23). Here, we observed low viral diversity within MHC-restricted CD8+ T cell epitopes immediately after CD8ɑ+ cell depletion in the aviremic PTCs (Figure 7), which is consistent with results from human PTCs described recently by Trémeaux et al. (41). Limiting viral replication, and by extent viral evolution, is one goal for HIV therapeutics aimed at treatment-free remission. Future studies that identify the immunological mechanism restricting viral replication among those harboring replication-competent proviruses could identify the essential features of an immunotherapy.

We do not fully understand how transient viremia following ART interruption impacts the composition of the rebound-competent viral reservoir in PTCs. Previous studies conducted in RM showed that while transient viral replication impacted the rebounding population following serial analytical treatment interruptions (ATI), this was limited and required a high degree of viral replication during the ATI (25). We utilized a similar logistic regression approach to determine whether the extent of pre-ART viral replication was a significant predictor of reactivation post-ART and whether reactivation post-ART or post-rechallenge influenced the likelihood of reactivation following depletion (Table 1). Across all animals and models evaluated, pre-ART viral load AUC was the predominant predictor of post-depletion reactivation; detection of a lineage during transient off-ART viremia was a significant predictor of post-depletion reactivation only in animal cy1037, whose viral population consisted primarily of the rechallenge virus. However, we must use caution when interpreting these results due to the low number of unique lineages reactivating during periods of off-ART viremia and the minimal viral replication observed during these periods. As such, while these results suggest that the rebounding viral population will stem from lineages comprising a large portion of the pre-ART population and that this population does not change significantly in the presence of low or undetectable viremia, we cannot definitively state that the reservoir cannot be reseeded or altered following a large burst of viral replication.

Following CD8ɑ+ cell depletion, we observed viral replication in all eight animals (Figure 1B). Sequencing the viral populations present prior to and following CD8ɑ+ cell depletion allowed us to examine the extent of sequence variation within CTL epitopes during the study (Figure 7). We found no evidence of immune escape within the three MHC-restricted CD8+ T cell epitopes evaluated in the aviremic animals at 7-10 days post depletion, but variation was common by ∼3-4 weeks post depletion (Figure 7). Importantly, given that rechallenge virus was only detected on the day of rechallenge (Supplemental Figure 2) and at a single time point in the aviremic animal with detectable rechallenge virus post-depletion (Figures 3 and 7), the presence of epitope sequences matching the inoculum observed in the plasma is unlikely to be derived from the rechallenge virus.

Detecting wild type SIV epitope sequences at the first post-depletion time point could be the result of two possibilities. First, there may have been limited viral evolution in the reservoir during PTC in these animals. Alternatively, viruses with wild type epitope sequences may have had a growth advantage over any variant viruses once the CD8ɑ+ cells were depleted. While we cannot distinguish between these possibilities with this data set, our results are in line with the hypothesis that early ART initiation and the presence of a small viral reservoir in MCM preserves wild type epitope sequences such that the CTL are more capable of virus suppression at the time of ART interruption. Based on the 3–4-week delay in variant detection observed here, we posit that SIV did not evolve during PTC in the aviremic animals because CD8+ cells were acting to suppress viral replication. One alternative hypothesis for the absence of immune escape variants despite CD8+ cell-mediated control is a non-cytotoxic CD8+ T cell mechanism that acts to suppress virus transcription without selecting for CTL escape variants (42–44). Future studies will need to compare viral variation from the reservoir during ART to that which is present after ART interruption in MCMs to characterize the extent to which antiviral CD8+ cells may be able to restrict viral replication and evolution during periods of PTC, potentially by employing next-generation sequencing technologies such as single-cell RNAseq or single genome amplification of viruses present in tissues.

Initiating ART during early HIV infection is the only intervention that is currently believed to increase the likelihood for PTC (3,45–47). We found that SIV-infected MCMs that began ART 8 wpi had a shorter time to viral rebound and a greater extent of pre- and post-ART viral sequence diversity within the CTL epitopes examined when compared to animals that initiated ART at 2 wpi (Figures 1, 6, 7, and Supplemental Figure 4). These results are consistent with previous studies reporting associations between later ART initiation, increased viral diversity, and shorter time to viral rebound following treatment interruption (48,49), as well as limited viral evolution and the maintenance of immune escape variants during ART treatment (6,50). Importantly, we determined that animals with later ART initiation had an increased number of viral variants present in the replicating viral population when ART was initiated. Therefore, our study supports the hypothesis that even a slight delay in ART initiation may allow for new variants to accumulate in the viral population, both prior to ART initiation and during ART treatment, reducing the possibility of establishing PTC.

Importantly, viruses isolated from the animals in the Late ART cohort had immune escape mutations in CTL epitopes examined prior to ART initiation, which may have impacted the ability of CD8+ T cells to suppress viral replication after ART interruption. To better understand the impact of immune escape on time to viral rebound following ART interruption, future studies could consider using a pre-escaped challenge virus in MCMs with defined MHC genetics. Pre-escaped viruses have previously been used to show impaired viral control during acute infection when compared to non-escaped, wild type virus (51), as well as inefficient recognition by CTL (38). When applied to future studies of PTC, the inclusion of similar pre-escaped viruses may allow researchers to evaluate the extent to which antiviral CD8+ T cell responses are required to establish and maintain PTC, as pre-existing CTL responses to the viral inoculum would be absent.

Unique to this study, we examined if the PTCs were susceptible to rechallenge with an isogenic viral strain by quantifying the proportion of virus containing a barcode, representing the initial challenge virus (SIVmac239M), and the proportion lacking a barcode, representing the rechallenge challenge virus (SIVmac239). Previous studies have shown that animals infected with an attenuated SIV strain or a Simian-human immunodeficiency virus (SHIV) strain were afforded protection from rechallenge with a similar, but genetically distinct, virus (52–55). Similar to these studies, we found that while most animals were protected from isogenic rechallenge, as determined by the low frequency of detectable rechallenge virus (SIVmac239) present in the plasma following depletion of CD8ɑ+ cells (Figure 3), this protection was highly variable, with one viremic animal exhibiting a viral population consisting of nearly 100% rechallenge virus following rechallenge and one aviremic animal containing only a single time point where the rechallenge virus was detectable post-depletion. While the proportion of rechallenge virus was low in these animals (Figure 3), it highlights that the rechallenge virus was able to establish a persistent, low-level infection despite immune control of plasma viremia, and that this rechallenge virus could be controlled spontaneously. Future studies examining differences in susceptibility to reinfection, particularly in PTCs, may have important implications in our understanding of reservoir dynamics.

This study does have certain limitations. Of note, the viremic and aviremic animals were part of a previous therapeutic vaccine study; however, no differences in viral kinetics were observed during the two-month period of follow up (31). Interestingly, the CD8+ T cells in animals that received the vaccine regimen (closed shapes) appeared to reappear in the plasma more rapidly following CD8ɑ+ cell depletion than animals that did not receive the vaccine regimen (open shapes) (Supplemental Figure 3A). Unfortunately, due to the heterogeneity observed in these animals, we are unable to draw conclusions regarding the relationship between the rate at which CD8ɑ+ cells repopulate the blood and the composition of the rebounding viral population.

Cumulatively, this study suggests a unique predisposition for MCM to establish PTC, highlights the importance of early ART initiation in restricting viral diversity and enhancing the likelihood of PTC, and emphasizes the importance of antiviral CD8+ cells in controlling viral replication during PTC. Of note, we propose the unique property of MCM to become PTCs. Further investigation into this potential model may allow us to explore the contribution of antiviral CD8+ cells to PTC. We provide evidence that while the composition of the viral reservoir remained stable during ART treatment and periods of PTC, small bursts of transient viral replication may influence this composition, even if viremia is spontaneously re-controlled. This finding highlights the importance of understanding mechanisms supporting viral reservoir maintenance in the absence of ongoing detectable viral replication. Further, we found that animals with a longer duration of SIV infection prior to ART initiation had more diverse viral populations pre- and post-ART, as well as detectable viremia substantially earlier than their Early ART counterparts (Figures 6 and 7). In sum, the data presented here provides evidence for a nonhuman primate model of PTC, which may allow researchers to more comprehensively understand the contribution of antiviral CD8+ cells in establishing and maintaining control of viremia in the absence of continual ART, as well as how these cells may act to restrict viral evolution during periods of PTC.

## Materials and Methods

### Animal care and use statement

All samples used in this study were collected from a previous study (31,32). A list of animal IDs, challenge virus, MHC genotype as determined by Wiseman et al (56), and cohort is provided in Supplemental Table 1.

### SIVmac239M infection and SIVmac239 rechallenge

The animal study was originally described in Harwood et. al. (31,32). For the Early ART cohort, eight MCM positive for at least one copy of the M3 haplotype were infected intravenously with 10,000 infectious units (IU) of SIVmac239M suspended in 1mL phosphate-buffered saline (PBS). Sixteen months following SIVmac239M infection, all animals were rechallenged intravenously with 100 TCID_50_ SIVmac239 suspended in 0.5mL. For the Late ART cohort, five MCM were infected intrarectally with 3,000 TCID50 of SIVmac239.

### CD8ɑ+ cell depletion

Eight SIVmac239M-infected MCM (Supplemental Table 1) were depleted of CD8ɑ+ cells as described in Harwood et al (31,32). Briefly, animals were given a single intravenous infusion of 50mg/kg CD8ɑ-depleting antibody (MT807R1). The rhesus macaque IgG1 recombinant Anti-CD8α (MT807R1) monoclonal antibody was engineered and produced by the Nonhuman Primate Reagent Resource (NIH Nonhuman Primate Reagent Resource Cat# PR-0817, RRID:AB_2716320). Depletion of CD8ɑ+ T cells was confirmed using a flow cytometry panel (32).

### SIV *gag* viral load quantification

SIV viral loads were quantified using a *gag* qPCR assay as previously described (57). Briefly, viral RNA was isolated from plasma using a Maxwell Viral Total Nucleic Acid kit (Promega), reverse transcribed, and amplified with the SuperScript III Platinum one-step quantitative RT-PCR system (Thermo Fisher Scientific). Viral RNA was quantified by quantitative PCR (qPCR) analysis on a LightCycler480 (Roche) and compared to an internal standard curve on each run. The assay used has a detection limit of 100 *gag* copy equivalents (ceq) per mL of plasma. The limit of detection (100 SIV *gag* ceq/mL) was reported when the viral load was at or below the limit of detection.

### SIV barcode sequencing

SIV barcode sequencing was performed as previously described (24) for all samples with a viral load of at least 1000 viral copies/mL of plasma or lymphoid tissue. Briefly, samples were reverse transcribed using SuperScriptIII (ThermoFisher Scientific) and a primer specific for a region adjacent to the barcode (Vpr.cDNA: 5’-CAG GTT GGC CGA TTC TGG AGT GGA TGC-3’ at position 6406–6380) per manufacturer’s instructions. Cycling conditions were as follows: 50°C for 1hr; 55°C for 1hr; 70°C for 15min; 10°C hold. 2U of RNAse H (ThermoFisher Scientific) was then added to the reaction, and samples were incubated at 37°C for 20min. Following cDNA synthesis, samples were PCR amplified using High Fidelity Platinum Taq (ThermoFisher Scientific) per manufacturer’s instructions and custom primers containing the Illumina index and adapter sequences (Supplemental Table 2). PCR cycling conditions were as follows: 94°C for 2min; 40x [94°C for 15sec, 60°C for 90sec, 68°C for 30]; 68°C for 5min; 10°C hold. Following PCR amplification, samples were subjected to a 1.2:1 AMPure bead clean-up step, then quantified using a Qubit High Sensitivity quantification assay (ThermoFisher Scientific) and amplicon lengths were determined using an Agilent BioAnalyzer (Agilent). Samples were then pooled equimolarly and sequenced on a 2×150 Illumina MiSeq.

### SIVmac239M barcode analysis

SIV samples were sequenced from plasma that had a minimum viral load 1000 copies/mL of plasma, corresponding to approximately 60 vRNA input templates. SIV barcode sequences were identified from the R1 read using a tool developed in the Keele lab and available at https://github.com/KeeleLab. All samples were sequenced in duplicate, and samples with less than 5000 reads were excluded. For each time point, barcode counts (the number of times a given barcode was identified in that sample) were pooled. SIVmac239M barcodes were included in the analysis if they were present in the pre-ART peak viral load and were above the minimum input threshold of 1/minimum number of input templates. Barcode sequences with a hamming distance of 1 from a known barcode were included in the analysis if they met the above criteria, otherwise barcodes containing multiple mismatches were discarded as sequencing artifacts.

### Whole-genome sequencing

SIV whole-genome sequencing was performed as previously described for samples with at least 1000 viral copies/mL of plasma (58). Briefly, vRNA was isolated from plasma using the Maxwell Viral Total Nucleic Acid kit (Promega). Four overlapping amplicons (A, B, C, and D) spanning the entire SIV coding sequence were generated for each sample using a Superscript III one-step reverse transcription (RT)-PCR system with high-fidelity Platinum Taq (Invitrogen) and the primers listed in Supplemental Table 3 with the following PCR cycling conditions: 50°C for 60min; 94°C for 2min; 2x [94°C for 15sec, 60°C for 1min, 68°C for 4min]; 2x [94°C for 15sec, 58°C for 1mion, 68°C for 4min]; 41x [94°C for 15sec, 55°C for 1min, 68°C for 4min]; 68°C for 10min; 10°C hold. Following PCR amplification, samples were subjected to a 1:1 AMPure bead clean-up step, then quantified using a Qubit High Sensitivity quantification assay (ThermoFisher Scientific). The amplicons for all samples were pooled equimolarly and used to generate uniquely tagged libraries using a Nextera XT kit (Illumina), per the manufacturer’s instructions. Samples were then sequenced in-house on a 2×250 Illumina MiSeq at 10pM with a 1% PhiX spike.

In the case of samples with amplicon dropouts and low viral loads, samples were amplified using an SIV multiplex PCR protocol as previously described (59). Briefly, cDNA was generated from vRNA using Superscript IV (Invitrogen). Samples were then amplified using the Q5 Polymerase (New England BioLabs) and the following PCR cycle conditions: 98°C for 30sec, 35x [95°C for 15sec and 65°C for 5 min]; 4°C hold. Amplified products were quantified using a Qubit dsDNA High Sensitivity assay (Thermo Fisher) and prepared for sequencing using an Illumina TruSeq kit per the manufacturer’s instructions. Prepared samples were pooled equimolarly and sequenced in-house on a 2×250 Illumina MiSeq at 10pM with a 10% PhiX spike.

### Epitope Identification and analysis

Following sequencing and demultiplexing, paired-end reads were merged using bbtools (60). Merged fastq files were mapped to the SIVmac239M reference and trimmed based on a minimum quality score. The epitope sequences for each animal and time point were identified, translated, and counted by amino acid sequence using a computational pipeline available on our github (https://github.com/rvmoriarty/PTC_MCM).

### Flow cytometry

To determine the amount and phenotype of CD8+ T cells present in the PBMC following CD8ɑ+ cell depletion was conducted as previously described (31,32). Briefly, previously cryopreserved PBMCs isolated from whole blood were used to assess the frequency of T cell populations longitudinally. Cells were thawed, washed with R10, and rested for 30 minutes at room temperature in a buffer consisting of 2% FBS in 1X PBS (2% FACS buffer) with 50nM dasatinib (Thermo Fisher Scientific). Cells were then washed with 2% FACS buffer with 50nM dasatinib and incubated at room temperature with Gag_386-394_GW9 and Nef_103-111_RM9 tetramers for 45 minutes. Cells were washed with 2% FACS buffer with 50nM dasatinib and incubated with the remaining surface markers for 20 minutes at room temperature. Cells were then washed with 2% FACS buffer, fixed for a minimum of 20 minutes with 2% paraformaldehyde, and acquired immediately using a FACS Symphony A3 (BD Biosciences). The data were analyzed with FlowJo software for Macintosh (BD Biosciences, version 10.8.0). Cell subpopulations were excluded from analysis when the parent population contained < 50 events.

### Sequence availability

All whole-genome viral sequences used are available on the Sequence Read Archive (SRA) under accession number SUB15753123.

### Computational and statistical analysis

All scripts used to generate and organize data are available upon request or on our GitHub (https://github.com/rvmoriarty/PTC_MCM). SIV barcodes were identified using a computational pipeline written in R, which can be downloaded from https://github.com/KeeleLab. Computational simulations to calculate the expected number of viral lineages that would be detected in the rebounding virus population post-CD8ɑ+ cell depletion, given the extent of pre-ART viral replication, was conducted in R as previously described (25). The R package DescTools was used to calculate viral load area under the curve (AUC). Logistic regressions were calculated using the glm function in R. For the single model evaluating post-ART rebound (Table 1, Model 1), the pre-ART viral load AUC was the sole explanatory variable. For Model 2, pre-ART viral load AUC was used as the sole explanatory variable. The next two models included the pre-ART viral load AUC and a binary indicator variable (detection of the linage post-ART or post-rechallenge) as separate covariates. Bray-Curtis dissimilarity index was calculated using the vegan R package (61). Statistical analysis was done using Prism 9 and R. Shapiro-Wilk tests were used to determine normality. For normally distributed data, t-tests were performed. For non-normally distributed data, Mann-Whitney U tests were performed.

## Acknowledgments

These studies were supported by NIH R01 AI108415. Research reported in this publication was supported by the National Institute of Allergy and Infectious Diseases of the National Institutes of Health under Award Number T32AI55397, and in part by the Office Of The Director, National Institutes of Health under Award Number P51OD011106 to the WNPRC, University of Wisconsin-Madison. This project has been funded in part with federal funds from the National Cancer Institute, National Institutes of Health, under Contract No. 75N91019D00024. The content of this publication does not necessarily reflect the views or policies of the Department of Health and Human Services, nor does mention of trade names, commercial products, or organizations imply endorsement by the U.S. Government. We thank the animal care staff at the Wisconsin National Primate Research Center at the University of Wisconsin-Madison for the excellent care of the animals housed at these facilities. The authors additionally thank Virology Services, a member of Research Services, at the WNPRC for performing SIV viral load assays. The content is solely the responsibility of the authors and does not necessarily represent the official views of the National Institutes of Health.

---

**Supplemental Table 1.** Animal IDs, challenge virus, MHC genotype, and cohort for all animals used in this study. Animals are color coded by cohort. MHC genotypes were determined as previously described (56).

| Animal ID | Initial Challenge Virus | Shape | MHC Genotype | Cohort |
| --- | --- | --- | --- | --- |
| cy1036 | SIVmac239M | closed square | M3/M5 | Early ART Aviremic |
| cy1039 | SIVmac239M | closed up triangle | M3/M3 | Early ART Aviremic |
| cy1044 | SIVmac239M | open up triangle | M2/M3 | Early ART Aviremic |
| cy1035 | SIVmac239M | closed circle | M3/M4 | Early ART Viremic |
| cy1037 | SIVmac239M | open circle | M3/recM3M4 | Early ART Viremic |
| cy1043 | SIVmac239M | closed down triangle | M2/recM3M4 | Early ART Viremic |
| cy1040 | SIVmac239M | open square | M3/M3 | Early ART Viremic |
| cy1045 | SIVmac239M | open down triangle | M3/M5 | Early ART Viremic |
| cy0719 | SIVmac239 | closed circle | M1, M3 | Late ART |
| cy0721 | SIVmac239 | closed square | recM1/M3, M5 | Late ART |
| cy0723 | SIVmac239 | closed up triangle | M4, recM3/M1 | Late ART |
| cy0724 | SIVmac239 | closed down triangle | M1, M4 | Late ART |
| cy0726 | SIVmac239 | open circle | M2, M3 | Late ART |

**Supplemental Table 2.**
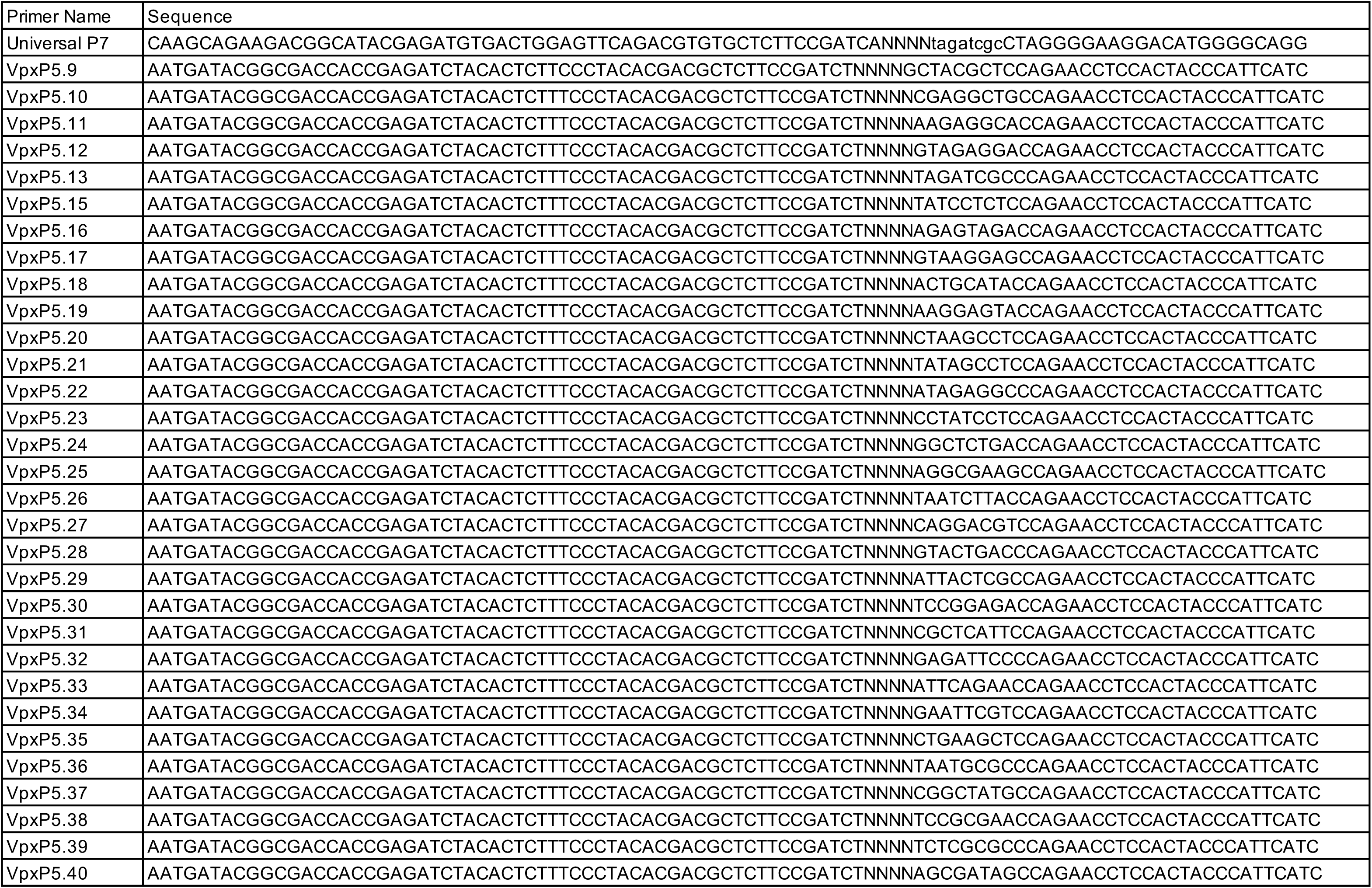
Primer sequences used in sequencing the SIVmac239M barcode.

**Supplemental Table 3.**
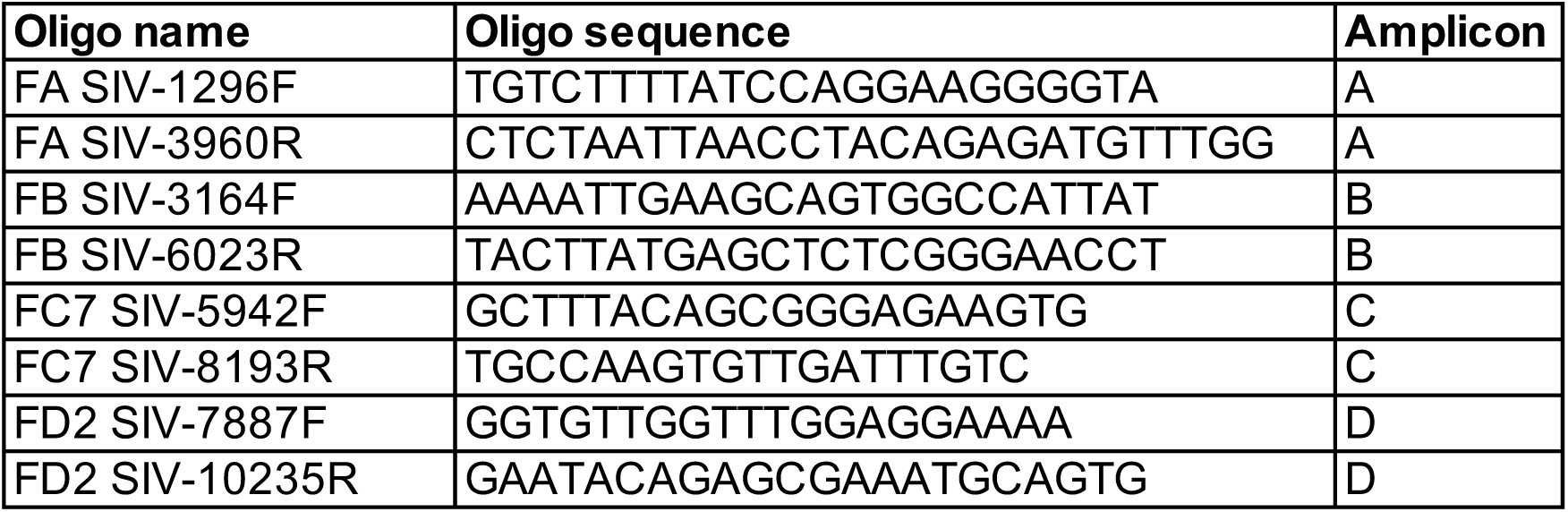
Primer names and sequences used for whole-genome sequencing using the four-amplicon approach, described in Sutton et al (58).

**Supplemental Figure 1.**
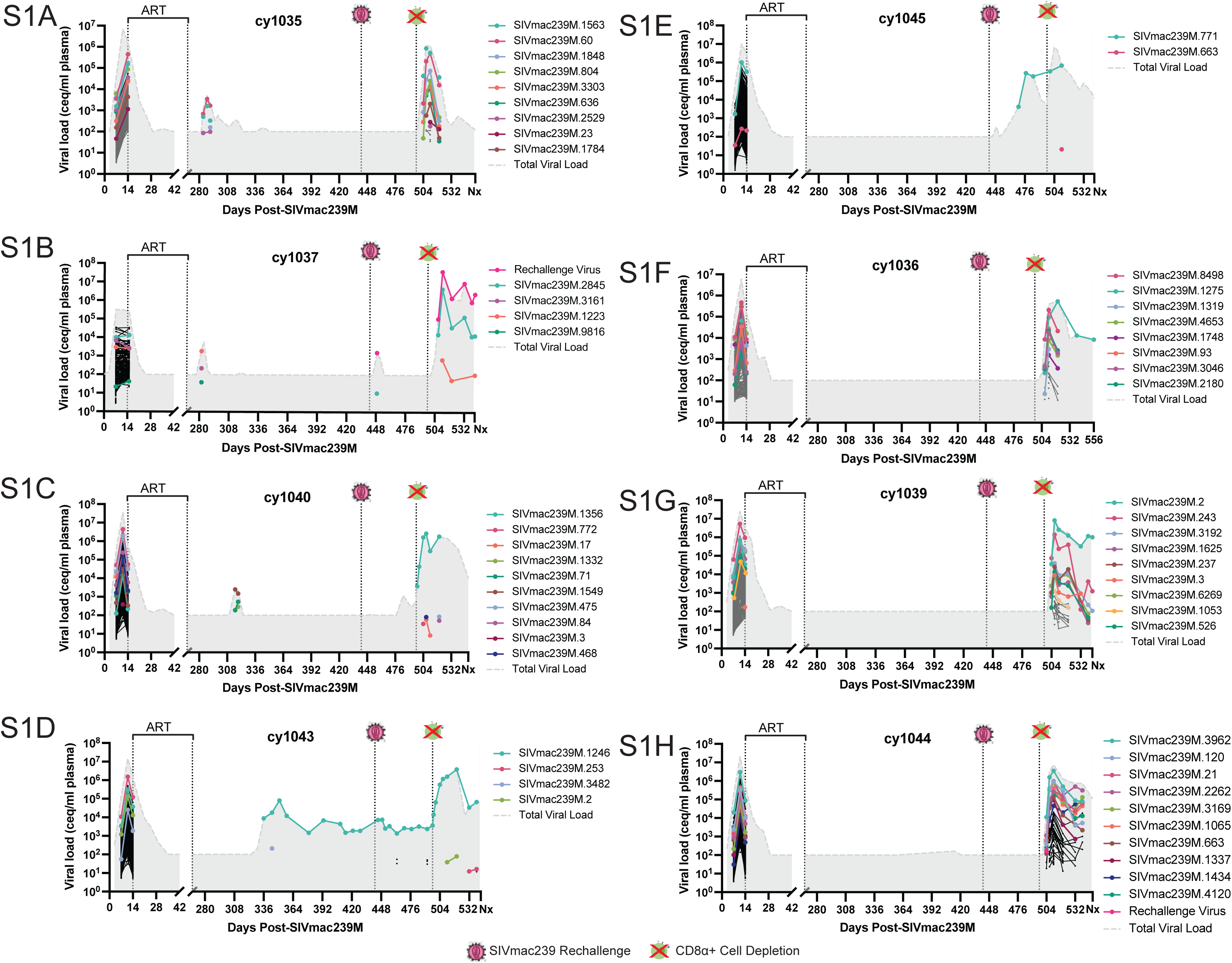
Log10 viral copies (Y axis) for each SIVmac239M lineage throughout the course of the study in viremic (A-E) and aviremic (F-H) animals. Each lineage is indicated by a different color, with only the top 10 lineages detected post-depletion colored due to color and space constraints. The remaining lineages are shown as grey lines and circles. Rechallenge virus is indicated in pink. Total log_10_ viral load is shown in the grey shaded area.

**Supplemental Figure 2.**
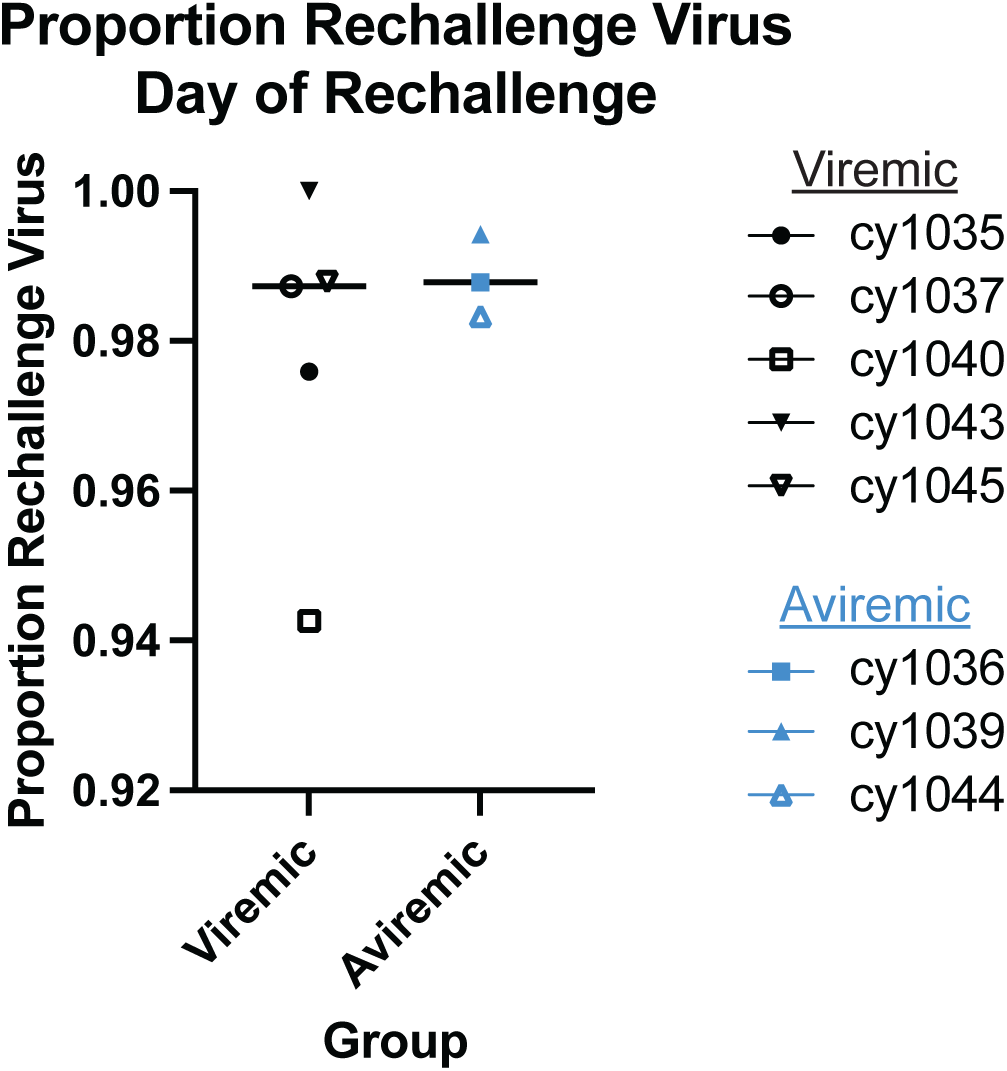
Proportion of rechallenge virus detected in the plasma immediately following intravenous infusion of SIVmac239. Each animal is indicated by a unique color and shape combination, with viremic animals shown in black and aviremic animals shown in blue. Line represents median of each group.

**Supplemental Figure 3.**
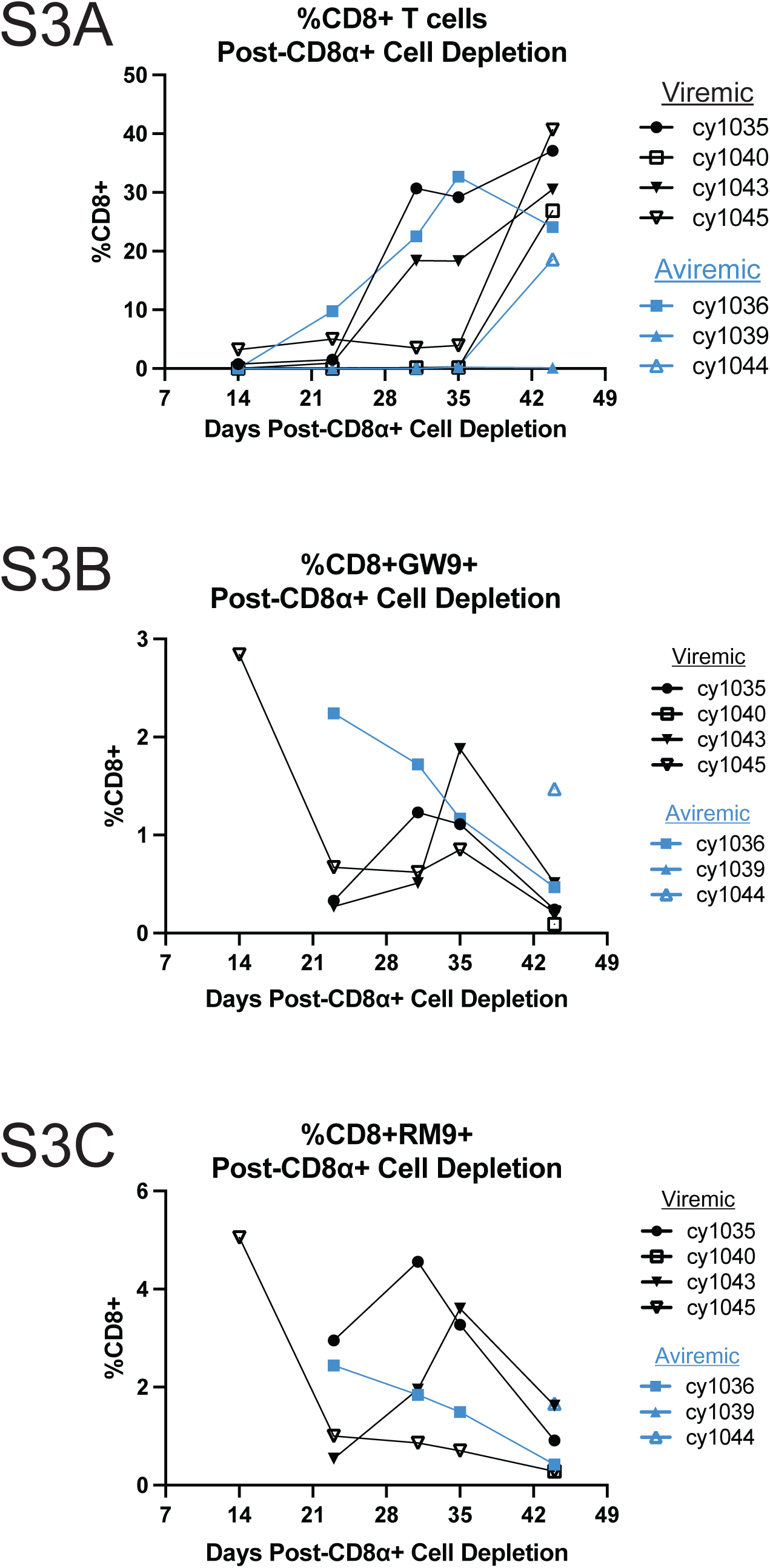
Frequency of bulk (A), Gag_386-398_GW9 tetramer+ (B), and Nef_103-111_RM9 tetramer+ (C) CD8+ T cells post-CD8ɑ+ cell depletion. Animals are coded by unique symbol and color combinations, with viremic animals in black and aviremic animals in blue. For subpopulation analysis, only samples in which the parent population of CD8+ cells had at least 50 events were included.

**Supplemental Figure 4.**
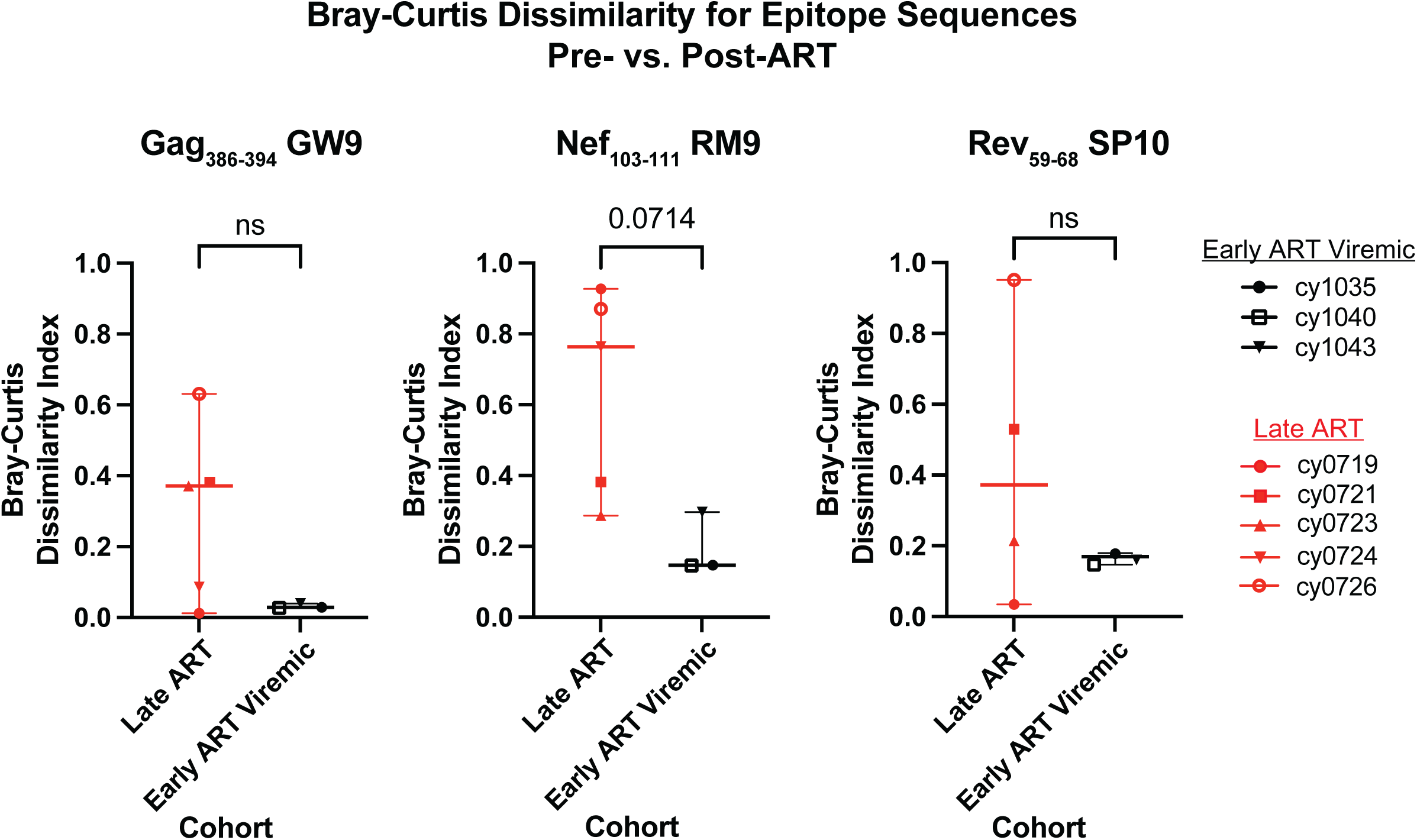
Bray-Curtis dissimilarity index for Gag_386-398_GW9 (left), Nef_103-111_RM9 (center), and Rev_59-68_SP10 (right) epitope sequences pre- and post-ART. Early ART viremic animals are in unique black symbols and Late ART animals are in unique red symbols. Significance was determined using Mann-Whitney U tests.

## References

1. Chun TW, Justement JS, Murray D, Hallahan CW, Maenza J, Collier AC, et al. Rebound of plasma viremia following cessation of antiretroviral therapy despite profoundly low levels of HIV reservoir: implications for eradication. Aids. 2010;24(18):2803–8.

2. Hocqueloux L, Prazuck T, Avettand-Fenoel V, Lafeuillade A, Cardon B, Viard JP, et al. Long-term immunovirologic control following antiretroviral therapy interruption in patients treated at the time of primary HIV-1 infection. Aids. 2010;24(10):1598–601.

3. Sáez-Cirión A, Bacchus C, Hocqueloux L, Avettand-Fenoel V, Girault I, Lecuroux C, et al. Post-Treatment HIV-1 Controllers with a Long-Term Virological Remission after the Interruption of Early Initiated Antiretroviral Therapy ANRS VISCONTI Study. Plos Pathog. 2013;9(3):e1003211.

4. Namazi G, Fajnzylber JM, Aga E, Bosch RJ, Acosta EP, Sharaf R, et al. The Control of HIV After Antiretroviral Medication Pause (CHAMP) Study: Posttreatment Controllers Identified From 14 Clinical Studies. J Infect Dis. 2018;218(12):1954–63.

5. Etemad B, Esmaeilzadeh E, Li JZ. Learning From the Exceptions: HIV Remission in Post-treatment Controllers. Front Immunol. 2019;10:1749.

6. Abdi B, Nguyen T, Brouillet S, Desire N, Sayon S, Wirden M, et al. No HIV-1 molecular evolution on long-term antiretroviral therapy initiated during primary HIV-1 infection. Aids. 2020;34(12):1745–53.

7. Bozzi G, Simonetti FR, Watters SA, Anderson EM, Gouzoulis M, Kearney MF, et al. No evidence of ongoing HIV replication or compartmentalization in tissues during combination antiretroviral therapy: Implications for HIV eradication. Sci Adv. 2019;5(9):eaav2045.

8. Kearney MF, Spindler J, Shao W, Yu S, Anderson EM, O’Shea A, et al. Lack of Detectable HIV-1 Molecular Evolution during Suppressive Antiretroviral Therapy. Plos Pathog. 2014;10(3):e1004010.

9. Josefsson L, Stockenstrom S von, Faria NR, Sinclair E, Bacchetti P, Killian M, et al. The HIV-1 reservoir in eight patients on long-term suppressive antiretroviral therapy is stable with few genetic changes over time. Proc National Acad Sci. 2013;110(51):E4987–96.

10. Nicolas A, Migraine J, Dutrieux J, Salmona M, Tauzin A, Hachiya A, et al. Genotypic and Phenotypic Diversity of the Replication-Competent HIV Reservoir in Treated Patients. Microbiol Spectr. 2022;10(4):e00784–22.

11. Brooks K, Jones BR, Dilernia DA, Wilkins DJ, Claiborne DT, McInally S, et al. HIV-1 variants are archived throughout infection and persist in the reservoir. Plos Pathog. 2020;16(6):e1008378.

12. Abrahams MR, Joseph SB, Garrett N, Tyers L, Moeser M, Archin N, et al. The replication-competent HIV-1 latent reservoir is primarily established near the time of therapy initiation. Sci Transl Med. 2019;11(513).

13. Brodin J, Zanini F, Thebo L, Lanz C, Bratt G, Neher RA, et al. Establishment and stability of the latent HIV-1 DNA reservoir. Elife. 2016;5:e18889.

14. Borrow P, Lewicki H, Wei X, Horwitz MS, Peffer N, Meyers H, et al. Antiviral pressure exerted by HIV-l-specific cytotoxic T lymphocytes (CTLs) during primary infection demonstrated by rapid selection of CTL escape virus. Nat Med. 1997;3(2):205–11.

15. O’Connor DH, McDermott AB, Krebs KC, Dodds EJ, Miller JE, Gonzalez EJ, et al. A Dominant Role for CD8 + -T-Lymphocyte Selection in Simian Immunodeficiency Virus Sequence Variation. J Virol. 2004;78(24):14012–22.

16. Emu B, Sinclair E, Hatano H, Ferre A, Shacklett B, Martin JN, et al. HLA Class I-Restricted T-Cell Responses May Contribute to the Control of Human Immunodeficiency Virus Infection, but Such Responses Are Not Always Necessary for Long-Term Virus Control. J Virol. 2008;82(11):5398–407.

17. Price DA, Goulder PJR, Klenerman P, Sewell AK, Easterbrook PJ, Troop M, et al. Positive selection of HIV-1 cytotoxic T lymphocyte escape variants during primary infection. Proc National Acad Sci. 1997;94(5):1890–5.

18. Miura T, Brumme CJ, Brockman MA, Brumme ZL, Pereyra F, Block BL, et al. HLA-Associated Viral Mutations Are Common in Human Immunodeficiency Virus Type 1 Elite Controllers. J Virol. 2009;83(7):3407–12.

19. Loffredo JT, Burwitz BJ, Rakasz EG, Spencer SP, Stephany JJ, Vela JPG, et al. The Antiviral Efficacy of Simian Immunodeficiency Virus-Specific CD8 + T Cells Is Unrelated to Epitope Specificity and Is Abrogated by Viral Escape. J Virol. 2007;81(6):2624–34.

20. Barouch DH, Kunstman J, Kuroda MJ, Schmitz JE, Santra S, Peyerl FW, et al. Eventual AIDS vaccine failure in a rhesus monkey by viral escape from cytotoxic T lymphocytes. Nature. 2002;415(6869):335–9.

21. Burwitz BJ, Giraldo-Vela JP, Reed J, Newman LP, Bean AT, Nimityongskul FA, et al. CD8+ and CD4+ cytotoxic T cell escape mutations precede breakthrough SIVmac239 viremia in an elite controller. Retrovirology. 2012;9(1):91.

22. Caetano DG, Côrtes FH, Bello G, Teixeira SLM, Hoagland B, Grinsztejn B, et al. Next-generation sequencing analyses of the emergence and maintenance of mutations in CTL epitopes in HIV controllers with differential viremia control. Retrovirology. 2018;15(1):62.

23. Loffredo JT, Friedrich TC, León EJ, Stephany JJ, Rodrigues DS, Spencer SP, et al. CD8+ T Cells from SIV Elite Controller Macaques Recognize Mamu-B*08-Bound Epitopes and Select for Widespread Viral Variation. Plos One. 2007;2(11):e1152.

24. Fennessey CM, Pinkevych M, Immonen TT, Reynaldi A, Venturi V, Nadella P, et al. Genetically-barcoded SIV facilitates enumeration of rebound variants and estimation of reactivation rates in nonhuman primates following interruption of suppressive antiretroviral therapy. Plos Pathog. 2017;13(5):e1006359.

25. Immonen TT, Fennessey CM, Lipkey L, Thorpe A, Prete GQD, Lifson JD, et al. Transient viral replication during analytical treatment interruptions in SIV infected macaques can alter the rebound-competent viral reservoir. Plos Pathog. 2021;17(6):e1009686.

26. Okoye AA, Duell DD, Fukazawa Y, Varco-Merth B, Marenco A, Behrens H, et al. CD8+ T cells fail to limit SIV reactivation following ART withdrawal until after viral amplification. J Clin Invest. 2021;131(8).

27. Pinkevych M, Fennessey CM, Cromer D, Tolstrup M, Søgaard OS, Rasmussen TA, et al. Estimating Initial Viral Levels during Simian Immunodeficiency Virus/Human Immunodeficiency Virus Reactivation from Latency. J Virol. 2018;92(2).

28. Swanstrom AE, Immonen TT, Oswald K, Pyle C, Thomas JA, Bosche WJ, et al. Antibody-mediated depletion of viral reservoirs is limited in SIV-infected macaques treated early with antiretroviral therapy. J Clin Invest. 2021;131(6).

29. Keele BF, Okoye AA, Fennessey CM, Varco-Merth B, Immonen TT, Kose E, et al. Early antiretroviral therapy in SIV-infected rhesus macaques reveals a multiphasic, saturable dynamic accumulation of the rebound competent viral reservoir. PLOS Pathog. 2024;20(4):e1012135.

30. Moriarty RV, Golfinos AE, Gellerup DD, Schweigert H, Mathiaparanam J, Balgeman AJ, et al. The mucosal barrier and anti-viral immune responses can eliminate portions of the viral population during transmission and early viral growth. Plos One. 2021;16(12):e0260010.

31. Harwood OE, Balgeman AJ, Weaver AJ, Ellis-Connell AL, Weiler AM, Erickson KN, et al. Transient T Cell Expansion, Activation, and Proliferation in Therapeutically Vaccinated Simian Immunodeficiency Virus-Positive Macaques Treated with N-803. J Virol. 2022;e01424–22.

32. Harwood OE, Matschke LM, Moriarty RV, Balgeman AJ, Weaver AJ, Ellis-Connell AL, et al. CD8+ cells and small viral reservoirs facilitate post-ART control of SIV replication in M3+ Mauritian cynomolgus macaques initiated on ART two weeks post-infection. PLOS Pathog. 2023;19(9):e1011676.

33. Budde ML, Greene JM, Chin EN, Ericsen AJ, Scarlotta M, Cain BT, et al. Specific CD8+ T Cell Responses Correlate with Control of Simian Immunodeficiency Virus Replication in Mauritian Cynomolgus Macaques. J Virol. 2012;86(14):7596–604.

34. Joseph SB, Swanstrom R, Kashuba ADM, Cohen MS. Bottlenecks in HIV-1 transmission: insights from the study of founder viruses. Nat Rev Microbiol. 2015;13(7):414–25.

35. Salazar-Gonzalez JF, Salazar MG, Keele BF, Learn GH, Giorgi EE, Li H, et al. Genetic identity, biological phenotype, and evolutionary pathways of transmitted/founder viruses in acute and early HIV-1 infection. J Exp Med. 2009;206(6):1273–89.

36. O’Connor SL, Becker EA, Weinfurter JT, Chin EN, Budde ML, Gostick E, et al. Conditional CD8+ T Cell Escape during Acute Simian Immunodeficiency Virus Infection. J Virol. 2011;86(1):605–9.

37. Harris M, Burns CM, Becker EA, Braasch AT, Gostick E, Johnson RC, et al. Acute-Phase CD8 T Cell Responses That Select for Escape Variants Are Needed to Control Live Attenuated Simian Immunodeficiency Virus. J Virol. 2013;87(16):9353–64.

38. Friedrich TC, McDermott AB, Reynolds MR, Piaskowski S, Fuenger S, Souza IP de, et al. Consequences of Cytotoxic T-Lymphocyte Escape: Common Escape Mutations in Simian Immunodeficiency Virus Are Poorly Recognized in Naïve Hosts. J Virol. 2004;78(18):10064–73.

39. Etemad B, Sun X, Li Y, Melberg M, Moisi D, Gottlieb R, et al. HIV post-treatment controllers have distinct immunological and virological features. Proc National Acad Sci. 2023;120(11):e2218960120.

40. Sharaf R, Lee GQ, Sun X, Etemad B, Aboukhater LM, Hu Z, et al. HIV-1 proviral landscapes distinguish posttreatment controllers from noncontrollers. J Clin Invest. 2018;128(9):4074–85.

41. Trémeaux P, Lemoine F, Mélard A, Gousset M, Boufassa F, Orr S, et al. In-Depth Characterization of Full-Length Archived Viral Genomes after Nine Years of Posttreatment HIV Control. Microbiol Spectr. 2023;11(1):e03267–22.

42. Walker CM, Erickson AL, Hsueh FC, Levy JA. Inhibition of human immunodeficiency virus replication in acutely infected CD4+ cells by CD8+ cells involves a noncytotoxic mechanism. J Virol. 1991;65(11):5921–7.

43. Zanoni M, Palesch D, Pinacchio C, Statzu M, Tharp GK, Paiardini M, et al. Innate, non-cytolytic CD8+ T cell-mediated suppression of HIV replication by MHC-independent inhibition of virus transcription. Plos Pathog. 2020;16(9):e1008821.

44. Mutascio S, Mota T, Franchitti L, Sharma AA, Willemse A, Bergstresser SN, et al. CD8+ T cells promote HIV latency by remodeling CD4+ T cell metabolism to enhance their survival, quiescence, and stemness. Immunity. 2023;

45. Maenza J, Tapia K, Holte S, Stekler JD, Stevens CE, Mullins JI, et al. How Often does Treatment of Primary HIV Lead to Post-Treatment Control? Antivir Ther. 2015;20(8):855–63.

46. Chéret A, Bacchus-Souffan C, Avettand-Fenoël V, Mélard A, Nembot G, Blanc C, et al. Combined ART started during acute HIV infection protects central memory CD4+ T cells and can induce remission. J Antimicrob Chemoth. 2015;70(7):2108–20.

47. Passaes C, Desjardins D, Chapel A, Monceaux V, Lemaitre J, Mélard A, et al. Early antiretroviral therapy favors post-treatment SIV control associated with the expansion of enhanced memory CD8+ T-cells. Nat Commun. 2024;15(1):178.

48. Li JZ, Aga E, Bosch RJ, Pilkinton M, Kroon E, MacLaren L, et al. Time to Viral Rebound After Interruption of Modern Antiretroviral Therapies. Clin Infect Dis. 2021;74(5):865–70.

49. Okoye AA, Hansen SG, Vaidya M, Fukazawa Y, Park H, Duell DM, et al. Early antiretroviral therapy limits SIV reservoir establishment to delay or prevent post-treatment viral rebound. Nat Med. 2018;24(9):1430–40.

50. Antar AAR, Jenike KM, Jang S, Rigau DN, Reeves DB, Hoh R, et al. Longitudinal study reveals HIV-1-infected CD4+ T cell dynamics during long-term antiretroviral therapy. J Clin Invest. 2020;130(7):3543–59.

51. Valentine LE, Loffredo JT, Bean AT, León EJ, MacNair CE, Beal DR, et al. Infection with “Escaped” Virus Variants Impairs Control of Simian Immunodeficiency Virus SIVmac239 Replication in Mamu-B*08 -Positive Macaques. J Virol. 2009;83(22):11514–27.

52. Reynolds MR, Weiler AM, Weisgrau KL, Piaskowski SM, Furlott JR, Weinfurter JT, et al. Macaques vaccinated with live-attenuated SIV control replication of heterologous virus. J Exp Medicine. 2008;205(11):2537–50.

53. Sharpe SA, Cope A, Dowall S, Berry N, Ham C, Heeney JL, et al. Macaques infected long-term with attenuated simian immunodeficiency virus (SIVmac) remain resistant to wild-type challenge, despite declining cytotoxic T lymphocyte responses to an immunodominant epitope. J Gen Virol. 2004;85(9):2591–602.

54. Berry N, Ham C, Mee ET, Rose NJ, Mattiuzzo G, Jenkins A, et al. Early Potent Protection against Heterologous SIVsmE660 Challenge Following Live Attenuated SIV Vaccination in Mauritian Cynomolgus Macaques. Plos One. 2011;6(8):e23092.

55. Yeh WW, Jaru-ampornpan P, Nevidomskyte D, Asmal M, Rao SS, Buzby AP, et al. Partial Protection of Simian Immunodeficiency Virus (SIV)-Infected Rhesus Monkeys against Superinfection with a Heterologous SIV Isolate. J Virol. 2009;83(6):2686–96.

56. Wiseman RW, Wojcechowskyj JA, Greene JM, Blasky AJ, Gopon T, Soma T, et al. Simian Immunodeficiency Virus SIVmac239 Infection of Major Histocompatibility Complex-Identical Cynomolgus Macaques from Mauritius. J Virol. 2007;81(1):349–61.

57. Cline AN, Bess JW, Piatak M Jr, Lifson JD. Highly sensitive SIV plasma viral load assay: practical considerations, realistic performance expectations, and application to reverse engineering of vaccines for AIDS. J Med Primatol. 2005;34(5-6):303–12.

58. Sutton MS, Ellis-Connell A, Moriarty RV, Balgeman AJ, Gellerup D, Barry G, et al. Acute-Phase CD4 + T Cell Responses Targeting Invariant Viral Regions Are Associated with Control of Live Attenuated Simian Immunodeficiency Virus. J Virol. 2018;92(21).

59. Moriarty RV, Fesser N, Sutton MS, Venturi V, Davenport MP, Schlub T, et al. Validation of multiplex PCR sequencing assay of SIV. Virol J. 2021;18(1):21.

60. Bushnell B. BBtools [Internet]. 2014. Available from: sourceforge.net/projects/bbmap/

61. Oksanen J, G S, F B, R K, P L, P M, et al. vegan: Community Ecology Program [Internet]. 2022. Available from: https://CRAN.R-project.org/package=vegan

